# Systematic Characterization of Thermal Stability Assay Parameters and Application in Discovery of Peptide-Protein Interactions

**DOI:** 10.64898/2026.05.06.723354

**Authors:** Daniel M. Richards, Fangyi (Coco) Zhai, Shaoxian Li, Qing Yu

## Abstract

Thermal proteome profiling (TPP) and its higher-throughput derivative, the proteome integral solubility alteration (PISA) assay, measure changes in protein thermal stability upon ligand binding or other perturbations and have been widely adopted in drug discovery and biomedical research. Though the PISA workflow is straightforward, key parameters, including detergent concentration, methods for removing denatured aggregates, and temperature range selection, vary across studies and can markedly influence assay outcomes. Yet these factors have not been systematically evaluated, limiting rational experimental design and data interpretation. Here, through a combined use of TPP, PISA, tandem mass tag (TMT)-based multiplexing, and computational simulation, we systematically characterize these parameters based on the melting behavior of ∼9,000 proteins. We find that reducing detergent concentration elevates apparent Tm by 1.5-2°C proteome-wide, and aggregate removal by filtration versus centrifugation further alters measurements. We leverage these observations to characterize how these parameters shape PISA, then apply selected conditions to identify the aminopeptidase NPEPPS as a previously uncharacterized binding partner of angiotensin II, a key vasoactive peptide hormone in blood pressure regulation. Together, this work provides a general framework for assay design and data interpretation, and extends the utility of PISA beyond small molecules to dissecting peptide-protein interactions, an increasingly important modality in drug discovery.

## INTRODUCTION

Ligand binding changes how proteins respond to thermal denaturation, a biophysical principle long exploited in differential scanning fluorimetry and related techniques (^1^). Martinez Molina et al. introduced the cellular thermal shift assay (CETSA), demonstrating that this principle could be applied to monitor target engagement directly in cells and tissues (^2^). Savitski et al. subsequently coupled thermal denaturation with quantitative mass spectrometry to enable proteome-scale target identification, termed Thermal Proteome Profiling (TPP) (^3^). In TPP, samples are heated across a temperature gradient, aggregated proteins removed, and remaining soluble protein quantified by tandem mass tag (TMT)-based proteomics. Proteins exhibiting altered thermal stability upon ligand treatment represent candidate targets. Beyond probing protein-ligand interaction, TPP has seen broader applications in biomedical research, including the assessment of protein complex architecture (^4^) and post-translational modification (^4, 5^).

However, TPP is inherently low throughput because it requires generating full thermal melt curves across many temperatures, making it tedious to scale for large screens. Gaetani et al. developed Proteome Integral Solubility Alteration (PISA), which increases throughput by pooling samples across the temperature gradient into a single measurement per condition (^6^). Rather than fitting individual melting curves, PISA quantifies the integral of soluble protein across temperatures, with ligand-induced stabilization manifesting as increased total soluble protein. Combined with TMT multiplexing, PISA substantially reduces instrument time while maintaining quantitative precision (^7, 8^).

Despite these advances, thermal shift methods show incomplete recovery of known ligand-protein interactions across studies, complicating data interpretation and limiting reproducibility (^7^). We attribute this in large part to the limited understanding of how various experimental conditions affect thermal stability measurements. The use of detergent at different concentrations, such as NP-40 from 0-0.5%, during cell lysis raises concerns for how this may impact ligand binding and solubility readout (^2, 3, 8, 9, 10^). Denatured aggregates are removed either by centrifugation or filtration, though their effects on the readout are unclear (^9, 11^). In addition, how these conditions together affect the selection of temperature ranges is less understood (^6, 12^). We therefore hypothesized that systematic evaluation of detergent concentration, aggregate removal strategy, and temperature range selection could improve detection sensitivity.

Peptide-protein interactions represent a particularly compelling yet underexplored application for thermal shift-based proteomics assays. Peptide hormones and neuropeptides regulate a wide range of essential physiological processes, including cardiovascular homeostasis, metabolism, immune signaling, and neural communication, by engaging diverse classes of intracellular and extracellular protein targets (^13, 14^). Dysregulation of peptide signaling pathways underlies numerous diseases, and peptides have emerged as an increasingly important therapeutic modality (^15^). However, systematic identification of peptide targets remains challenging. PISA is well suited to detect these interactions in native cellular environments at scale. Establishing well-characterized PISA workflows and the understanding of how assay conditions influence protein melting and solubility behavior is therefore essential to fully leverage this approach for studying peptide hormone biology and advancing peptide-based drug discovery.

Here, we characterize how the three aforementioned parameters can affect protein melting behavior in TPP, and consequently measurements in PISA. We measured ∼9,000 proteins and generated >6,000 high-quality melting curves (sigmoidal fits, R²>0.95) at two NP-40 concentrations. We compared filtration and centrifugation for aggregate removal. Both parameters shift apparent melting temperature (Tm) globally, with DNA and RNA binding proteins exhibiting particular sensitivity to the method used for aggregate removal which could be rescued by removal of DNA and RNA with benzonase. We further extended the prior analysis of temperature range selection, underscoring the balance between quantitative sensitivity and the depth of quantifiable proteome using PISA. We then applied PISA to profile angiotensin II (AngII) interactors, an octapeptide hormone essential in regulating blood pressure and fluid balance, identifying the puromycin-sensitive aminopeptidase NPEPPS as a novel AngII-binding protein and validating the inhibitory interaction biochemically.

## METHODS

### Cell culture and lysate preparation

HCT116 cells (ATCC Cat# CCL-247) were cultured in Dulbecco’s Modified Eagle’s Medium (HyClone™, SH30243.02) supplemented with 10% fetal bovine serum (FBS) and 1% penicillin-streptomycin at 37°C in a humidified atmosphere containing 5% CO2. Cells were grown to ∼70% confluence before harvesting, washed twice with ice-cold PBS, and harvested by scraping.

To prepare lysate, either fresh or frozen cell pellets were resuspended in lysis buffer consisting of PBS without Ca^2+^ and Mg^2+^ (HyClone™, SH30028.02) with supplemented with either 0.5% or 0.1% NP-40 no protease inhibitors. Cells were lysed using probe sonication in 5-second on/off cycles followed by 20 strokes with a 21G needle on a 5 mL syringe. All harvest and lysis procedures were performed on ice. Crude lysate was centrifuged at 650× g for 2 minutes at 4°C to verify complete lysis and remove large cell debris. Protein concentration was determined by BCA assay (Thermo Fisher, Cat #23225).

### Thermal Proteome Profiling (TPP)

Lysates were diluted to 1 mg/mL total protein concentration and 120 µL was aliquoted into 8-strip PCR tubes. Samples were heated using a thermal cycler across a regularly spaced temperature gradient from 48°C to 65°C (^3, 7, 16^). The total ramp from room temperature and hold time was 3 minutes, followed by a 1-minute decrease to 23°C with a 5-second hold before transfer to ice for ∼5 minutes.

After heating, denatured protein aggregates were removed by one of two methods: (a) filtration through 0.45 μm PTFE filter plates, pre-washed with 100 µL lysis buffer, followed by centrifugation at 3,000× g for 10 minutes, with quality control verification that equal volumes were recovered across all samples; or (b) centrifugation at 20,000× g for 60 minutes at 4°C, collecting the supernatant and immediately transferring to new tubes. Soluble protein recovery was quantified by BCA assay for each temperature point. For TPP experiments using benzonase, lysate was treated at 1 U/µL after heating for 20 minutes at RT before removing aggregate.

An equal volume was taken from each sample, with protein mass ranging from ∼36 µg in the lowest temperature channels to ∼8 µg in the highest temperature channels, and mixed 1:1 with 2× sample prep buffer (400 mM EPPS, 10 mM TCEP, 4% SDS) for 15 minutes with shaking. Iodoacetamide was added to a final concentration of 10 mM and incubated at room temperature in the dark for 20 minutes. DTT was then added to a final concentration of 10 mM to quench excess iodoacetamide for 10 minutes with shaking before proceeding to SP3-based cleanup, protein digestion, TMT labeling, and mass spectrometry acquisition(^17, 18^).

Melting curves were fitted using the TPP R package as described (^11^). Proteins were classified as non-melting if the fitted curve had an R^2^ below 0.8 and a bottom plateau of 0.4 or higher(^16^).

### Proteome Integral Solubility Alteration (PISA)

For PISA experiments (^6, 8, 12^), lysates were prepared as described above and incubated with angiotensin II [MCE, Cat# HY-13948]) or vehicle control (0.05% DMSO) at room temperature for 60 minutes. Cell lysate (120 µL at 1 mg/mL) was then distributed among 4 PCR tubes. Samples were heated at each specified temperature range with identical cycle times to TPP experiments (^12^) then pooled. Aggregates were removed by centrifugation at 20,000× g for 60 minutes at 4°C (filtration data not shown). Protein concentration was measured by BCA assay. An equal volume equivalent to ∼20 µg protein was taken from each sample and mixed 1:1 with 2× sample prep buffer (400 mM EPPS, 10 mM TCEP, 4% SDS) for 15 minutes with shaking. Iodoacetamide was added to a final concentration of 10 mM (20 minutes, RT, dark), followed by DTT to a final concentration of 10 mM (10 minutes) to quench iodoacetamide before proceeding to SP3-based cleanup.

### Solid-phase protein precipitation and sample preparation for mass spectrometry

SP3 bead cleanup was performed as described (^17^). After reduction and alkylation, proteins were incubated in a 1:1 mixture of hydrophilic and hydrophobic Sera-Mag SpeedBeads (Cytiva, GE45152105050350 & 65152105050350) in 60% ethanol for 15 minutes at room temperature with shaking. Beads were precipitated on a magnetic rack and washed three times with 80% ethanol, then resuspended in 20 µL of 200 mM EPPS pH 8.5 containing LysC (Fujifilm, Cat# 121-05063) at a 1:100 enzyme:substrate ratio. Proteins were digested overnight at room temperature with shaking, followed by addition of 10 µL of 200 mM EPPS pH 8.5 containing trypsin (Thermo Fisher, Cat# 90057) at 1:50 enzyme:substrate ratio for a further 6 hours at 37°C with shaking.

Peptides were labeled with TMTpro 18-plex or 35-plex reagents (Thermo Fisher, Cat# A52045; A40000817; A40000818) (^18, 19^). Briefly, for 20 µg protein digest in ∼30 µL EPPS buffer, 11 µL of acetonitrile and 4 µL TMT (10 mg/mL) were added. Peptides were labeled for 1 hour at room temperature with shaking. Reactions were quenched by adding 5% hydroxylamine to a final concentration of 0.3% for 15 minutes at room temperature. Labeled samples were pooled, acetonitrile was removed by vacuum centrifugation, and peptides were desalted using a 50 mg Sep-Pak (Waters, WAT054955).

### Basic pH reversed-phase fractionation

Pooled TMT-labeled peptides equivalent to ∼160 µg from each experiment were fractionated using an Agilent 1260 HPLC system equipped with an Agilent ZORBAX 2.1 mm × 250 mm C18 column with 3.5 µM pore size. Peptides were separated in 10 mM ammonium bicarbonate over 45 minutes at a flow rate of 0.225 mL/min. A total of 96 fractions were collected and concatenated into 24 pools. Fractions were desalted by using C18 disk (Empore, 98-0604-0218-1) before LC/MS analysis(^12, 17, 18^).

### Mass spectrometry acquisition

Orbitrap Eclipse (SPS-RTS-MS3). Fractionated samples were analyzed on an Orbitrap Eclipse Tribrid mass spectrometer (Thermo Fisher) coupled to a Vanquish Neo UHPLC system (^20^). Peptides were separated on a 100 µm × 30 cm column packed with 2.6 µm C18 particles using an 85-minute gradient from 8% to 30% Buffer B (95% acetonitrile in 0.125% formic acid). MS1 spectra were acquired in the Orbitrap at 60,000 resolution (at m/z 200) with an AGC target of 4 × 105 and maximum injection time of 50 ms. MS2 spectra were acquired in the ion trap using CID fragmentation (normalized collision energy 35%) with an AGC target of 1 × 104 and maximum injection time of 50 ms. MS3 spectra were acquired using synchronous precursor selection (SPS) of the 10 most intense MS2 fragment ions, HCD fragmentation (normalized collision energy 55%), and detection in the Orbitrap at 50,000 resolution with an AGC target of 1 × 105 and maximum injection time of 120 ms.

Real-time search (RTS) was enabled to trigger MS3 scans only for confidently identified peptides (^20^).

### Data analysis

Database search and quantification. Raw files were searched using the Comet search engine (^21^) against the UniProt human proteome database. Search parameters included: trypsin digestion (cleavage after K/R) with up to 2 missed cleavages, 50 ppm precursor ion tolerance, and 1.0005 Da fragment ion tolerance for ion trap CID MS2 spectra. TMTpro (+304.2071 Da) on lysine residues and peptide N-termini and carbamidomethylation of cysteine (+57.0215 Da) were set as fixed modifications. Oxidation of methionine (+15.9949 Da) and N-terminal acetylation (+42.0106 Da) were set as variable modifications. Data were filtered to a 1% protein-level false discovery rate.

TMT reporter ion signal-to-noise ratios were extracted from SPS-MS3 spectra using a 0.002 Da window around the theoretical m/z of each TMTpro reporter ion (126 through 135n, 18 channels). Reporter ions were filtered at signal-to-noise ≥ 10 per channel. Data were normalized by equalizing the summed reporter ion intensities across all channels within each experiment.

TPP analysis. Melting curves were fitted using the TPP R package (^11^). The median apparent Tm and interquartile range were calculated for proteins meeting quality thresholds (R^2^ > 0.8).

PISA analysis. Log2 fold changes were calculated as the ratio of ligand-treated to vehicle control reporter ion signal-to-noise ratios. Statistical significance was assessed using Student’s t-test with Benjamini-Hochberg (BH) FDR correction where stated. Proteins were considered hits if they met thresholds of |log2FC| > 0.3 and BH-corrected q-value < 0.05. The small fold change threshold is selected consistent with established PISA studies as PISA generates compressed fold-change distribution of integral-solubility readouts (^6, 8, 12^). For the angiotensin II screen we applied a q-value cutoff in the main-figure comparisons (**Figure 5**), while the broader hit sets based on raw p-values are shown in **Supplementary Figure 4**.

PISA fold-change simulation. To relate TPP-derived Tm values to the PISA readout, we simulated 10,000 paired melting curves. In each iteration, a control Tm was drawn from the experimental Tm distribution (0.5% NP-40, centrifugation), and a ΔTm was drawn from a uniform distribution U (−6, +6)°C. Control and ligand-shifted sigmoidal curves were simulated as previously described (^12^) by calculating the intensity at a given temperature as:

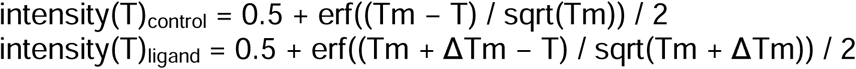

where erf is the error function, and sqrt is the square root function. The simulated intensity(*T*)_control_ or intensity(*T*)_ligand_ corresponds to an intensity normalized to the intensity at the top plateau of the melting curve. The sum of intensity (*T*)_control_ and the sum of intensity(*T*)_ligand_ values for selected temperature ranges were used to calculate Sm_control_ (i.e., ∑ intensity(*T*)_control_), Sm_ligand_ (i.e., ∑ intensity(*T*)_ligand_), and fold change (i.e., Sm_ligand_/Sm_control_).

Gene Ontology enrichment. Gene Ontology enrichment analysis was performed using the clusterProfiler R package (^22^) with all detected proteins used as the background set and Fisher’s exact test with Benjamini-Hochberg FDR correction.

Effect size analysis. Cohen’s d effect sizes were calculated to compare aggregate removal methods and detergent concentrations (^23^):

d = (M1 – M2) / s_pooled_

where M1 and M2 are the group means and spooled is the pooled standard deviation. Effect magnitudes were interpreted using established thresholds: |d| < 0.2 negligible, < 0.5 small, < 0.5 medium, ≥ 0.8 large.

Known angiotensin-pathway interactors and processing enzymes spanning angiotensin I–IV were compiled into a reference set (AGTR1, AGTR2, MAS1, AGT, REN, ACE, ACE2, MME, CMA1, ENPEP, ANPEP, LNPEP, PRCP, PREP, THOP1, NLN, DPP3, DPP4, CTSA, CTSG, CTSD, CPA3, PITRM1) against which detected proteins were cross-referenced and those detected in our dataset are labeled in red on figure 5, serving as our true positive set for AngII.

### AlphaFold structure prediction and structural feature analysis

AlphaFold2 structure predictions were obtained for all proteins in the dataset with fitted melting curves (R2 > 0.8) and Tm values from TPP data. For proteins with multiple isoforms, the canonical UniProt sequence was used (^24^). Structural features were calculated using Bio3D (^25^) and PyMOL (Schrödinger, LLC) as described below.

The predicted Local Distance Difference Test (pLDDT) score was extracted for each residue from B-factor fields of the AlphaFold model. The geometric mean pLDDT across all N residues was calculated as a metric of overall folding confidence:

geometric mean pLDDT = exp((1/N) × Σi ln(pLDDTi)) where pLDDTi > 0 for all residues i.

The fraction of disordered residues was defined as the proportion with pLDDT < 70 (^26^). Maximum protein dimension was calculated as the greatest pairwise Cα–Cα distance in the predicted structure.

Intramolecular contact density was defined as the number of Cα–Cα contacts per residue, where a contact is counted when the interatomic distance is < 8.0 Å and the sequence separation is ≥ 4 residues:

contact density = N_contacts_ / N_residues_, where contact between residues i and j is defined as: ||r_i_ – r_j_|| < 8.0 Å and |i – j| ≥ 4

Relative contact order was calculated as (^27^):

CO = (1 / (N_contacts_ × L)) × Σ ΔS_ij_

where ΔS_ij_ = |i – j| is the sequence separation between contacting residues, N_contacts_ is the total number of contacts, and L is protein length.

For NPEPPS–angiotensin II interaction modeling, AlphaFold 3 cofolding predictions were generated with angiotensin II (sequence: DRVYIHPF) and full-length human NPEPPS (^28^). Contact analysis between angiotensin II residues and NPEPPS was performed using PyMOL and Maestro.

### NPEPPS activity assay

Recombinant human NPEPPS was obtained from R&D Systems (Cat# 6410-ZN-010) (^29^). Aminopeptidase activity was measured using Leu-7-amino-4-methylcoumarin (Leu-MCA; Cayman Chemical, Cat# 62480-44-8) as substrate. Assays were performed in 96-well black plates in buffer containing 25 mM HEPES and 1 mM DTT at pH 7 with final concentrations of 0.2 ng/µL NPEPPS and 10 µM Leu-MCA. Fluorescence (excitation 380 nm, emission 460 nm) was monitored at one-minute intervals for 5 minutes at room temperature using a plate reader. Curve shown is from the 5 minute timepoint.

For IC_50_ determination, NPEPPS was pre-incubated with varying concentrations of angiotensin II (0– 70 µM) for 15 minutes at room temperature before addition of substrate. Initial reaction velocities were calculated from the linear phase of fluorescence increase. IC50 values were determined by fitting the data to a four-parameter logistic equation using GraphPad Prism (GraphPad Software, San Diego, CA).

## RESULTS

### Detergent concentration and aggregate removal method alter melting curves and apparent Tm

To investigate how detergent concentration and strategy to remove denatured aggregates impact TPP and PISA readout, we first performed TPP on HCT116 cell lysates prepared in PBS containing either 0.5% or 0.1% NP-40 (Figure 1A). Lysates were heated from 48°C to 65°C. Soluble fractions were collected by either 0.45 μm PTFE filtration or centrifugation (20,000 × g, 60 min, 4°C)(^7^). Protein recovery ranged from approximately 36 μg (60% input) at the lowest temperature to 8 μg (13.3% input) at the highest temperature per TMT channel. Each detergent condition occupied one 16-plex within a 32-plex experiment(^19^). Following high-pH reversed-phase fractionation into 24 pools, samples were analyzed on an Orbitrap Eclipse using SPS-RTS-MS3 acquisition (^20^).

**Figure 1.**
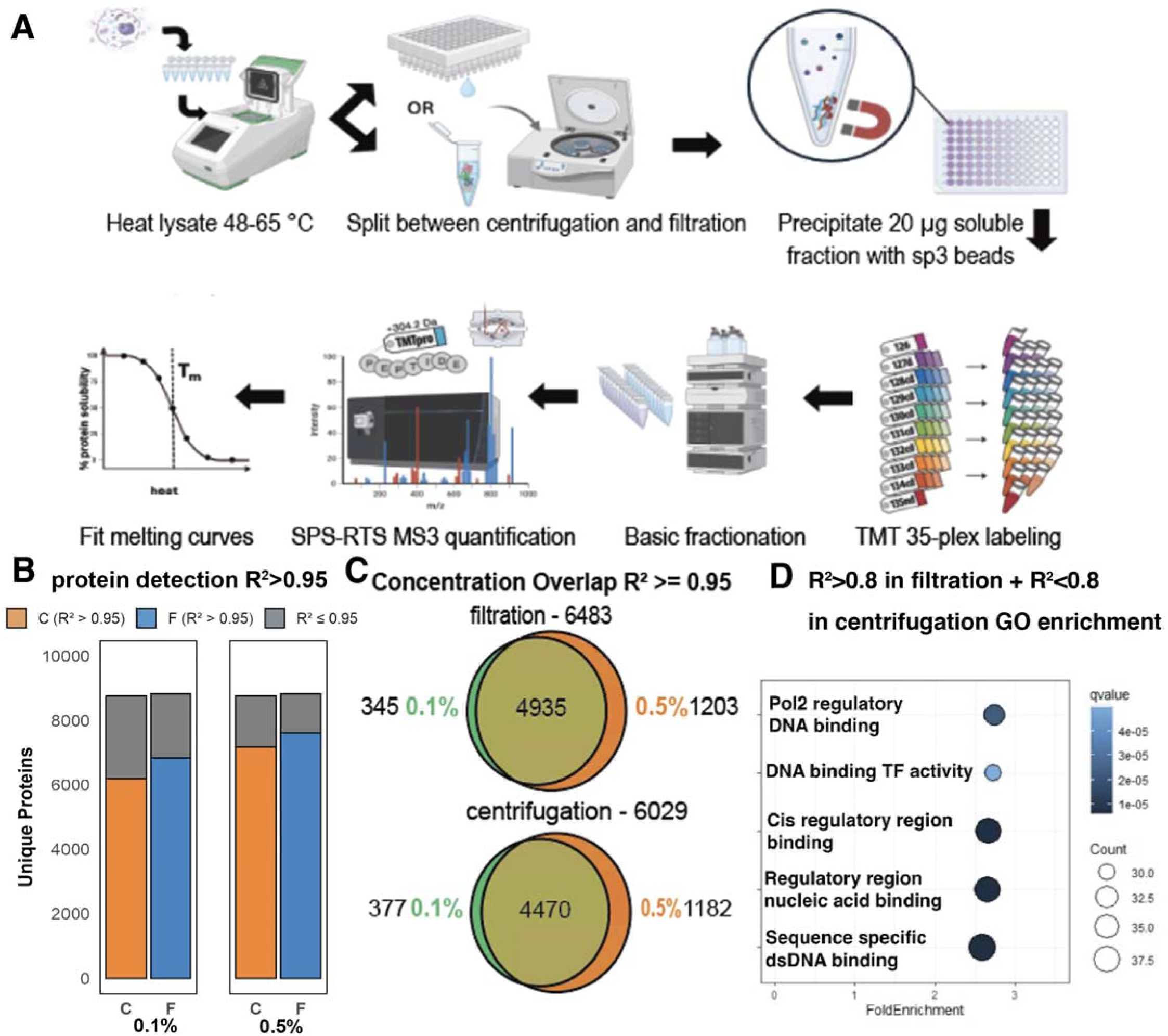
Thermal Proteome Profiling to compare detergent concentration and solubilization method impact on observed Tm. (A) TPP workflow: 1. heat across a gradient 2. collect soluble fraction 3. precipitate protein and lysC/trypsin digest, 4. TMT label, offline fractionation, 5. MS quantification 6. Fit melting curves (B) Number of proteins detected and number of proteins that fit a melting curve with R^2^>0.95 in two detergent concentrations. Higher detergent and filtration conditions have more proteins that fit a curve. C = centrifugation, F = filtration. (C) Venn diagram of proteins with good melting curves in 0.1% and 0.5% NP-40 conditions. (D) Proteins that are a good fit (R^2^>0.95) in filtration and a bad fit (R^2^<0.8) in centrifugation conditions are enriched in DNA binding activity.

Protein melting curves were fit using the TPP R package (^11^). In total, 8852 proteins were quantified. All four conditions (two detergent concentrations × two aggregate removal methods) yielded >34,000 curves with R² ≥ 0.95 (**Figure 1B**). Using higher detergent concentration and filtration for lysate solubilization increased the number of high-quality curve fits (Figure 1C). Using 0.5% NP-40 yielded 1,748 additional high-quality sigmoidal curves (13.4% increase) compared to 0.1% NP-40, while aggregate removal by filtration produced 1,084 (8.1%) more high-quality curves than centrifugation.

Gene Ontology analysis of proteins showing high quality fits with filtration (R^2^>0.8) but poor fits with centrifugation (R^2^<0.8) revealed enrichment for DNA-binding annotations **(Figure 1D)**. MACROH2A1 exemplifies this category **(Supplementary Figure 1D)**. This pattern suggests that DNA-binding proteins are particularly sensitive to the choice of aggregate removal strategy. Treating lysate with benzonase after heating was sufficient to rescue well-fit melting curves for several DNA-binding proteins, including MACROH2A1 **(Supplementary Figure 1D)**, indicating that bound DNA can mask or distort the apparent melting behavior of these proteins. Proteins that only produced well-fit curves with benzonase were significantly enriched only for double-stranded DNA-binding. Conversely, proteins that only fit curves in the absence of benzonase were primarily enriched for ribosomal proteins, potentially reflecting a dependence of ribosomal protein complex integrity on intact rRNA.

Examination of mean bottom plateau values across all curves confirmed that filtration removes aggregated protein more completely than centrifugation **(Supplementary Figure 1A)**. Proteins fitting curves only with centrifugation showed no functional enrichment.

### Filtration reduces and compresses observed Tm distributions

Filtration decreased median observed Tm by approximately 1°C relative to centrifugation (**Figure 2A**). Scatter plots directly comparing Tm values between aggregate removal methods at matched detergent concentrations confirmed this systematic shift (**Figure 2D**). Bottom plateau values in fitted curves were lower with filtration at both detergent concentrations (Supplementary Figure 1A), consistent with more efficient aggregate clearance. We performed Cohen’s d effect size analysis with Benjamini-Hochberg FDR-corrected Wilcoxon tests to estimate the contributions of aggregate removal strategy and detergent concentration to melting-curve parameters, interpreting effect magnitudes using established thresholds (|d| < 0.2 negligible, < 0.5 small, < 0.5 medium, ≥ 0.8 large)(^23^). Aggregate removal method exerted a larger effect on curve slope than detergent concentration, with centrifugation yielding steeper transitions overall **(Figure 2F, Supplementary Figure 1B)**. The interquartile range of apparent Tm was narrower with filtration than centrifugation. Inflection point, representing the temperature of maximum curve steepness, was impacted most by changes to aggregate removal method compared to detergent, and a similar pattern was seen for slope **(Figure 2E, Supplementary Figure 1C)**. Given that Tm denotes 50% protein loss while inflection point marks maximum slope, differential effects on these parameters indicate that filtration and centrifugation produce curves of distinct shape rather than simply shifted curves, likely driven by the systematic decrease in plateau for filtration.

**Figure 2.**
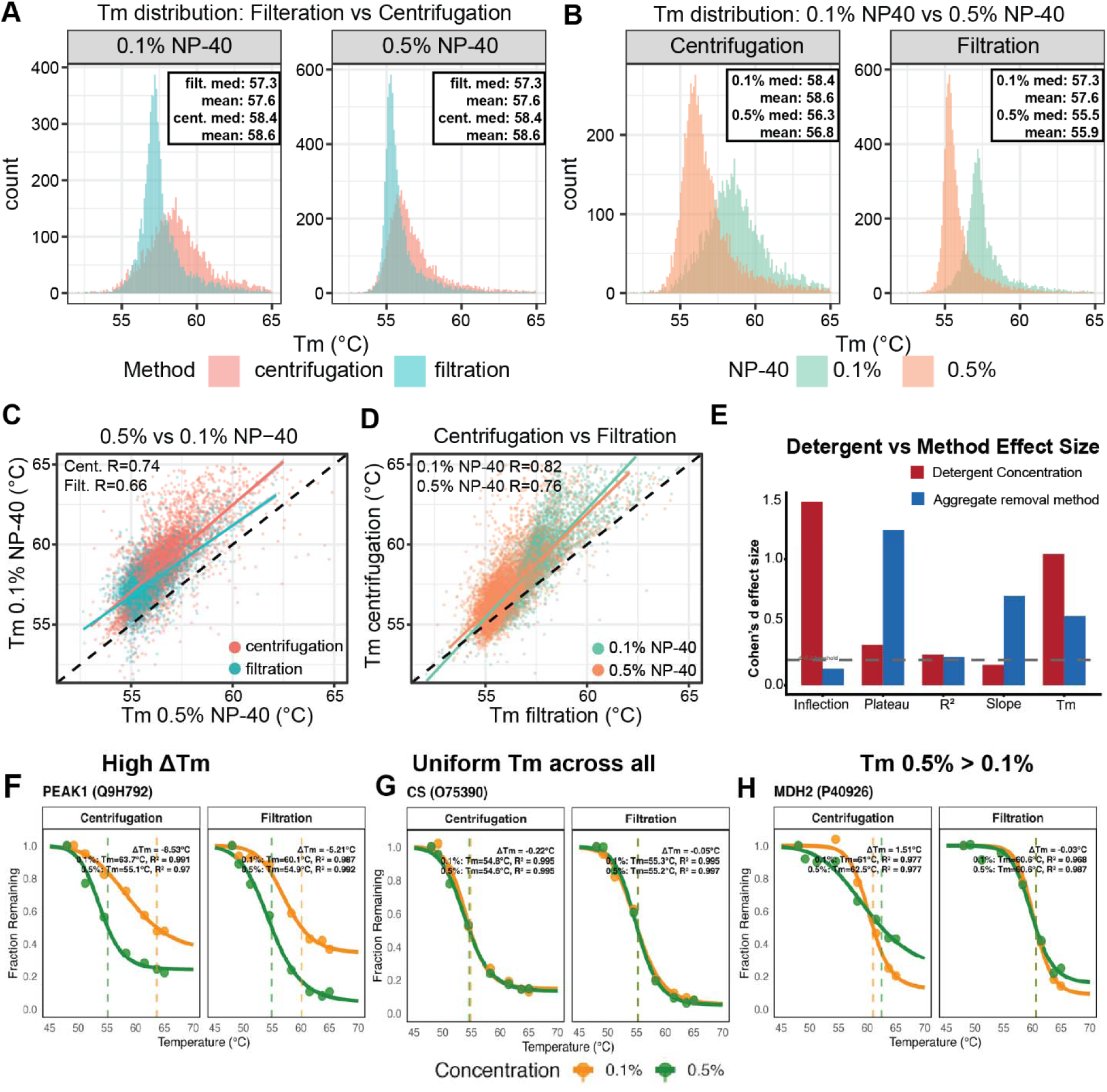
Impact of detergent concentration and aggregate removal strategy on observed Tm. (A) Distribution of observed Tm for 0.1% and 0.5% NP-40 conditions with solubilization methods overlapped. Filtration on average reduces observed Tm. (B) Distribution of observed Tm for centrifugation and filtration with detergent concentrations overlapped. Reducing detergent concentration increases observed Tm. (C) Scatterplot overlap of Tm values for 0.1% and 0.5% NP-40 conditions. Dotted diagonal line represents y=x. (D) Scatterplot overlap of Tm values for centrifugation and filtration methods. Dotted diagonal line represents y=x. (E) Effect size (Cohens d) for detergent or aggregate removal method foreach of the melting curve parameters given by the TPP R package: Inflection point, plateau, R^2^, slope and Tm. (F) Representative melting curve for PEAK1, a protein with high delta Tm between 0.1% and 0.5% NP-40 (G) Representative melting curve for CS, a protein with similar Tm between 0.1% and 0.5% NP-40 and aggregate removal method (H) Representative melting curve for MDH2, a protein which had an observed Tm higher in 0.5% than 0.1%.

Even though filtration yielded promising data, the filtration plates we used occasionally exhibited inconsistent flow rates, limiting its suitability for large scale screening. Therefore, subsequent PISA experiments employed centrifugation exclusively. For filtration-based workflows, a post-aggregate-removal BCA across all channels is an essential quality-control step. Because PISA uses an equal volume of soluble fraction from all samples that came from the same starting input, uneven recovery propagates directly into the fold-change estimate. For a successful assay, we target even recovery of 30-40% of the starting protein to remove enough denatured proteins to resolve binding events while preserving proteome depth.

### Reduced detergent concentration elevates observed Tm

Observed Tm distributions at both detergent concentrations are shown in **Figure 2B**. Scatter plots comparing Tm between detergent concentrations with matched aggregate removal methods demonstrated the shift **(Figure 2C)**. Detergent concentration had a larger effect magnitude on apparent Tm than aggregate removal method **(Figure 2E)**. 0.1% NP-40 lysates exhibited mean Tm values 1.5-2°C higher than 0.5% NP-40 lysates, with the precise difference depending on aggregate removal method and whether mean or median was compared **(Figure 2A**, **Figure 2B)**. This is consistent with Reinhard et al. who compared 0.4% NP-40 to no detergent and observed a global decrease in the Tm of ATP binding proteins (^10^). Individual proteins exhibited variable responses **(Figure 2F)**. Although most followed global trends, a subset showed minimal difference between conditions **(Figure 2G)**, and others displayed inverted patterns **(Figure 2H)**.

### Detergent sensitivity correlates with predicted structural order

It has previously been established that protein length is negatively correlated with Tm (^16, 30, 31^). To investigate if detergent sensitivity, defined as the magnitude of ΔTm between 0.5% and 0.1% NP-40, was due to any characteristic structural features, we obtained AlphaFold structure predictions for all proteins detected in the TPP dataset (^24, 28^). Using a quartile ranking system similar to that described by Jarzab et al. 2020 (^16^), but applied to detergent sensitivity (|ΔTm|) rather than Tm, we observed that the top 25% of proteins with the highest |ΔTm| have the longest length **(Supplementary Figure 2A, Supplementary Figure 2B)**. We reasoned also that proteins which have more compact structure would have less room for detergent molecules to intercalate and change conformation, rendering them less sensitive. Indeed, |ΔTm| showed a significant negative correlation with average intramolecular contact density (centrifugation: ρ = 0.37; filtration: ρ = 0.43) and the geometric mean AlphaFold pLDDT (predicted Local Distance Difference Test) score (centrifugation: ρ = 0.36; filtration: ρ = 0.37) which is a proxy for folding confidence (^24^) **(Figure 3A, Supplementary Figure 2D)**.

**Figure 3.**
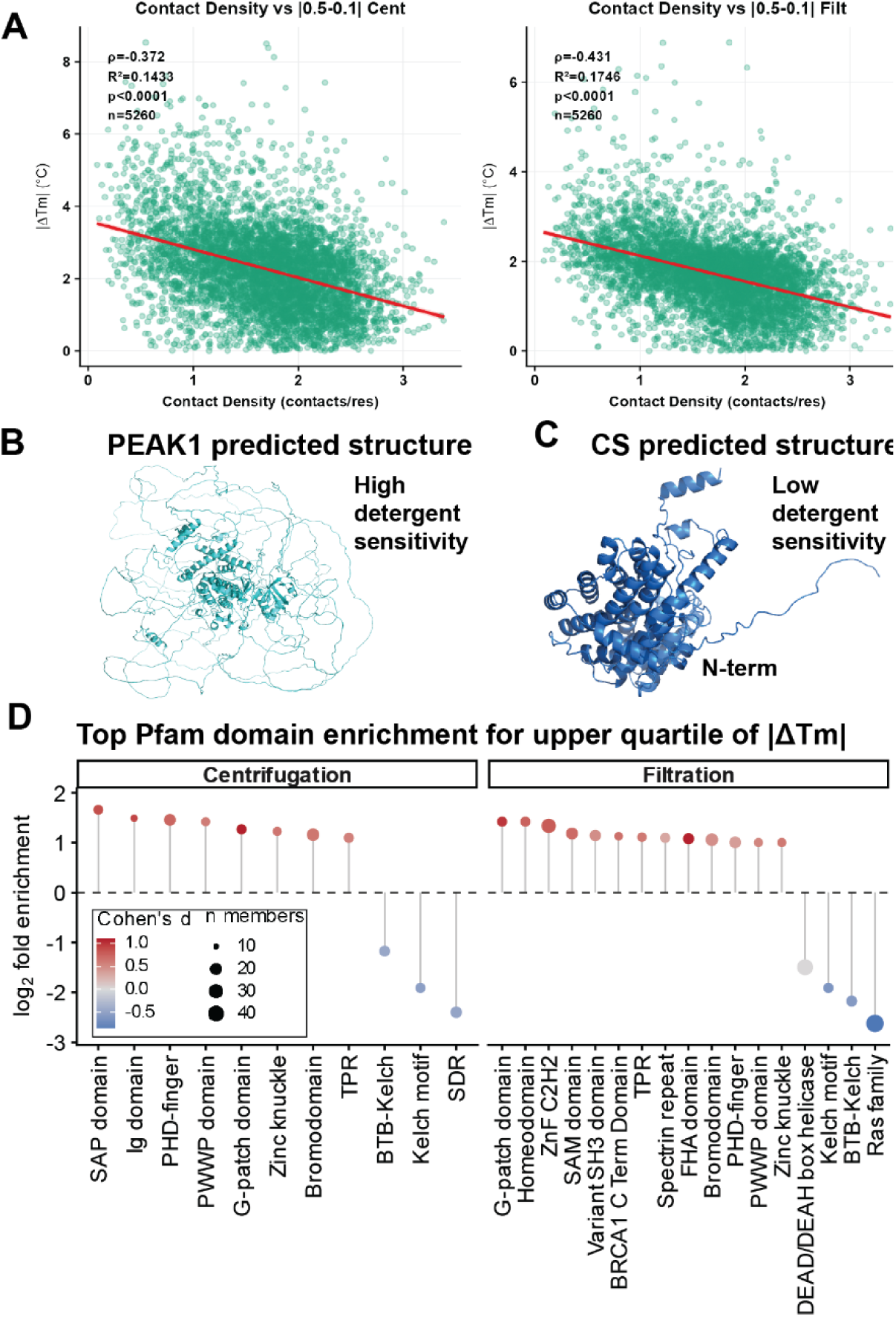
Detergent sensitivity correlates with structural features. (A) Contact density per residue of a protein assessed via pymol vs |ΔTm| (0.5% - 0.1%) for centrifugation (left) and filtration (right) methods. (B) Alphafold predicted structure of inactive tyrosine-protein kinase PEAK1, a protein with high detergent sensitivity that has many large, disordered loops (C) Alphafold predicted structure of citrate synthetase CS, a protein with no detergent sensitivity has a small compact structure (D) Pfam domains showing the strongest positive and negative enrichment among the most detergent-sensitive proteins (upper quartile of |ΔTm|), relative to the full proteome. Dots are colored by Cohen’s d effect size. Terms with fold enrichment>2 are shown.

Similarly, a modest correlation existed with maximum protein dimension (centrifugation: ρ = 0.31; filtration: ρ = 0.32) **(Supplementary Figure 2E)**. Notably contact density, pLDDT and max dimensions all had greater Spearman correlations compared to protein length (centrifugation: ρ = 0.17; filtration: ρ = 0.14) **(Supplementary Figure 2C)**. Examining a broader set of structural features reinforced this picture: across both aggregate-removal methods, |ΔTm| correlated positively with disorder, length, and molecular extension (radius of gyration, maximum dimension), and negatively with compactness and folding-confidence metrics (contact density, packing density, contact order, pLDDT). Melting and non-melting proteins separated along these same axes, with non-melting proteins showing greater disorder **(Supplementary Figure 3A, B)**. These results suggest that extended, disordered protein structure, rather than protein length alone, is the stronger determinant of susceptibility to detergent denaturation. Given the complexity inherent in analyzing predicted structures, we were nonetheless struck that experimental thermal behavior reflected defining structural features. Example predicted structures for PEAK1, a protein with high |ΔTm|= 8.53, show large disordered regions **(Figure 3B)**, compared to CS, a protein with low |ΔTm|= 0.22, where the only predicted disordered region is the N-terminal mitochondrial targeting sequence **(Figure 3C)**.

Consideration of predicted structure may therefore inform condition selection in targeted applications to maximize fold change and detectability of a protein of interest and clarifying false negative interactions for control compounds to better benchmark assay success. The choice of detergent concentration should balance the risk of disrupting detergent-sensitive proteins versus the need to sufficiently solubilize a larger fraction of the proteome for discovery work. When parsing the |ΔTm| detergent sensitivity metric by Pfam domains, we observe enrichment of several DNA/RNA binding domains (e.g., SAP domain, PHD-finger, PWWP, G-patch domain) among the most detergent-sensitive proteins **(Figure 3D)**. We also note that many DNA/chromatin-binding proteins are characteristically modular, containing substantial intrinsically disordered regions flanking the annotated domain, which may represent a major contributing factor. In addition, domains involved in protein-protein interactions (e.g., ankyrin repeat, tetratricopeptide repeat) are enriched and proteins possessing metabolic enzyme domains (e.g., short chain dehydrogenase, aldehyde dehydrogenase) are among the least detergent-sensitive **(Figure 3D)**.

### Simulation analysis of PISA temperature selection

Previous works by Gaetani et al. and Li et al. reported that narrowing the temperature range to the region of the most significant solubility changes, or to higher temperatures, could improve the dynamic range and fold change (FC) of the PISA readout (^6, 12^). However, increasing temperature to enhance FC comes at the expense of proteome coverage, as greater fractions of the proteins with lower Tm are lost. Using the experimentally determined Tm values from TPP obtained under 0.5% NP-40 and centrifugation-based aggregate removal, we simulated the relationship between FC in PISA and the remaining soluble fraction of the starting material. The median Tm determined by TPP was 56°C, prompting us to first simulate FC and proteome loss using a narrow 57-59°C temperature window. We simulated 10,000 paired curves, each consisting of a baseline melting curve with Tm randomly drawn from the experimental population and a shifted curve with a randomly sampled ΔTm between −6 and +6°C, mimicking destabilization and stabilization, respectively. For each pair, fold changes were calculated as ratios of areas under the curves (AUCs), an approximate for the FC in PISA. These simulations revealed a clear trend toward higher FC when the baseline Tm fell below the lower boundary of the PISA temperature range. For stabilized curves, the median log₂ FC was 0.71 (∼1.64-fold) for proteins with Tm < 57°C, compared with 0.40 (∼1.32-fold) for proteins with Tm ≥ 57°C (**Figure 4A**). Conversely, higher temperatures resulted in substantial loss of soluble material. For proteins with Tm < 57°C, only ∼45% of the total AUC remained soluble and thus measurable, compared to ∼63% for proteins with Tm ≥ 57°C. This effect was even more pronounced for destabilizing shifts, for which the median remaining soluble fraction dropped to ∼25% for proteins with Tm < 57°C (**Figure 4B**).

**Figure 4.**
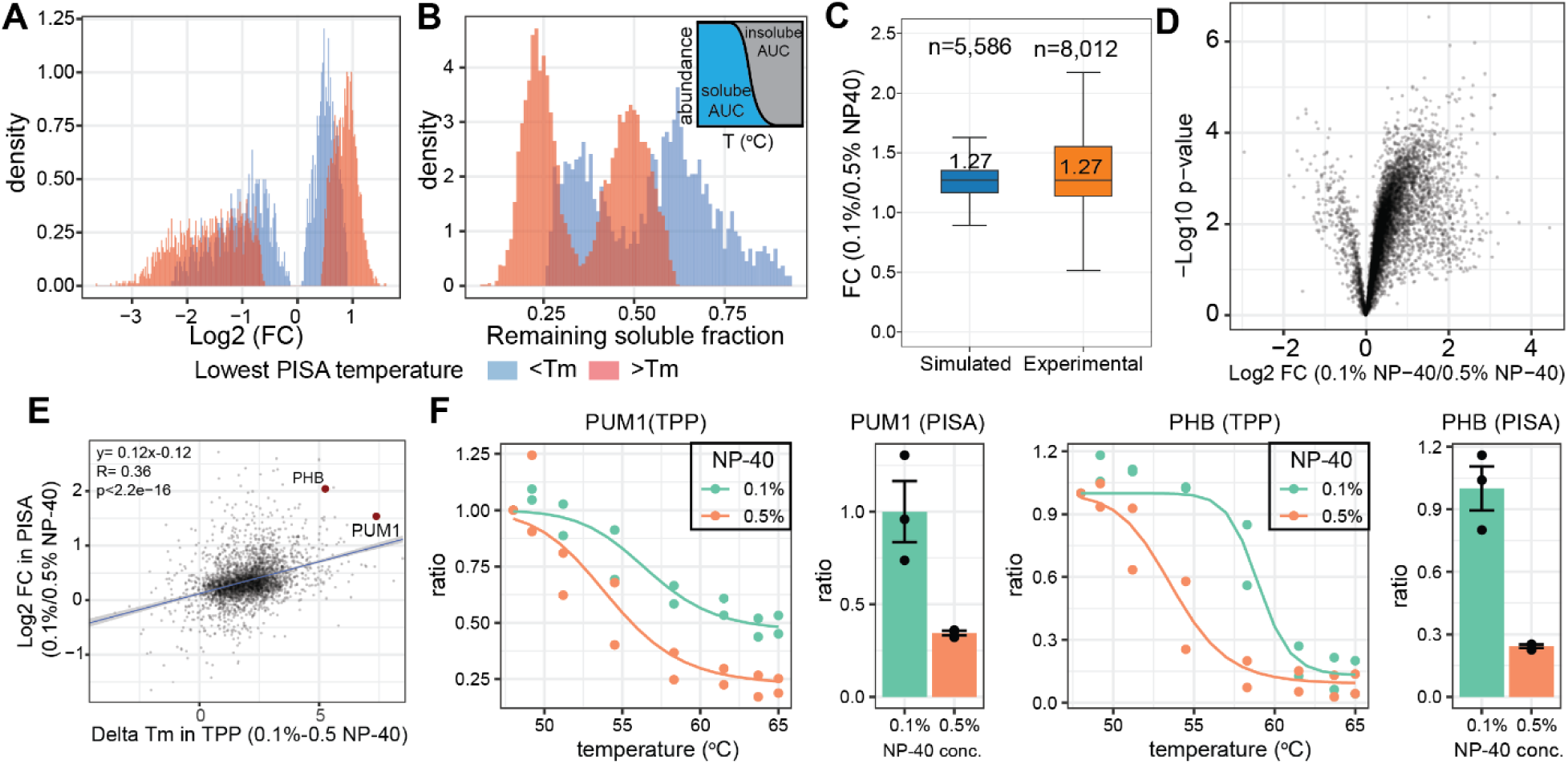
Balancing PISA temperature range selection for quantitative and qualitative sensitivity. (A) PISA simulation using random Tm values derived from TPP data. Heating above the Tm of a protein results in larger fold changes in PISA. (B) PISA simulation using random Tm values derived from TPP data. Heating above the Tm of a protein results in a greater fraction of the protein becoming insoluble and therefore removed from the final sample. (C) Simulated versus experimentally recovered protein abundance (n = 5,586 for simulated results; n = 8,012 for experimental data). The remaining soluble fraction of each protein was simulated and experimentally measured under 0.1% and 0.5% NP-40 conditions, respectively, and ratios were calculated for each protein. (D) Volcano plot comparing protein abundance in PISA under 0.1% and 0.5% NP 40 conditions. Majority of the proteome have higher abundance with lower NP-40 concentration. (E) A moderate correlation (R=0.36) between Log2 FCs from PISA data and ΔTm values from TPP comparing protein thermal stability using 0.1% and 0.5% NP 40. (F) TPP curves and PISA protein abundance of example proteins.

Similarly, our TPP data showed that higher NP-40 concentrations globally lowered Tm values, and as a result we anticipated greater protein loss when applying the same PISA temperature range under 0.5% NP-40 versus 0.1% NP-40. Indeed, aggregating AUCs across fitted melting curves across the 53-59°C range revealed that 0.1% NP-40 retained ∼27% more total soluble proteome relative to 0.5% NP-40. To test this simulation, we performed a PISA assay on vehicle-treated HCT116 lysate comparing 0.1% and 0.5% NP-40. The measured soluble protein abundance closely matched our estimates, showing a median 27% higher abundance at the lower detergent concentration (**Figure 4C, D**). Importantly, the fractional protein loss was not uniform, reflecting protein specific differences in Tm values and Tm shifts between the two detergent conditions. We observed a moderate correlation between Log2 FC (0.1%/0.5% NP-40) and ΔTm **(Figure 4E)**. For example, PHB exhibited Tm of 59.3 °C under 0.1% NP-40 but 54.0 °C under 0.5% NP-40. As a result, we observed a 4x decrease in soluble PHB abundance in PISA **(Figure 4F)**. These findings underscore the importance of jointly tuning detergent concentration and temperature to balance quantitative and qualitative sensitivity, particularly when specific proteins or protein classes are of interest.

### Angiotensin II exhibits detergent-dependent hit profiles

To assess whether PISA can be applied to discovery of peptide-protein interactions, we performed PISA with angiotensin II (AngII), an octapeptide hormone central to blood pressure regulation. The experiment was done under both 0.1% and 0.5% NP-40 lysis conditions to further assess detergent impact. Overall, 5 proteins were detected across conditions using a fold change threshold of log2(FC) > |0.3| and q-value < 0.05. 0.5% NP-40 yielded 2 more proteins passing the significance threshold than 0.1% NP-40 **(Figure 5A, B)**. Three proteins reached significance in both detergent conditions **(Figure 5A, B)**, including documented AngII-interacting peptidases dipeptidyl peptidase 3 (DPP3) and neurolysin (NLN) (^32, 33^).

**Figure 5.**
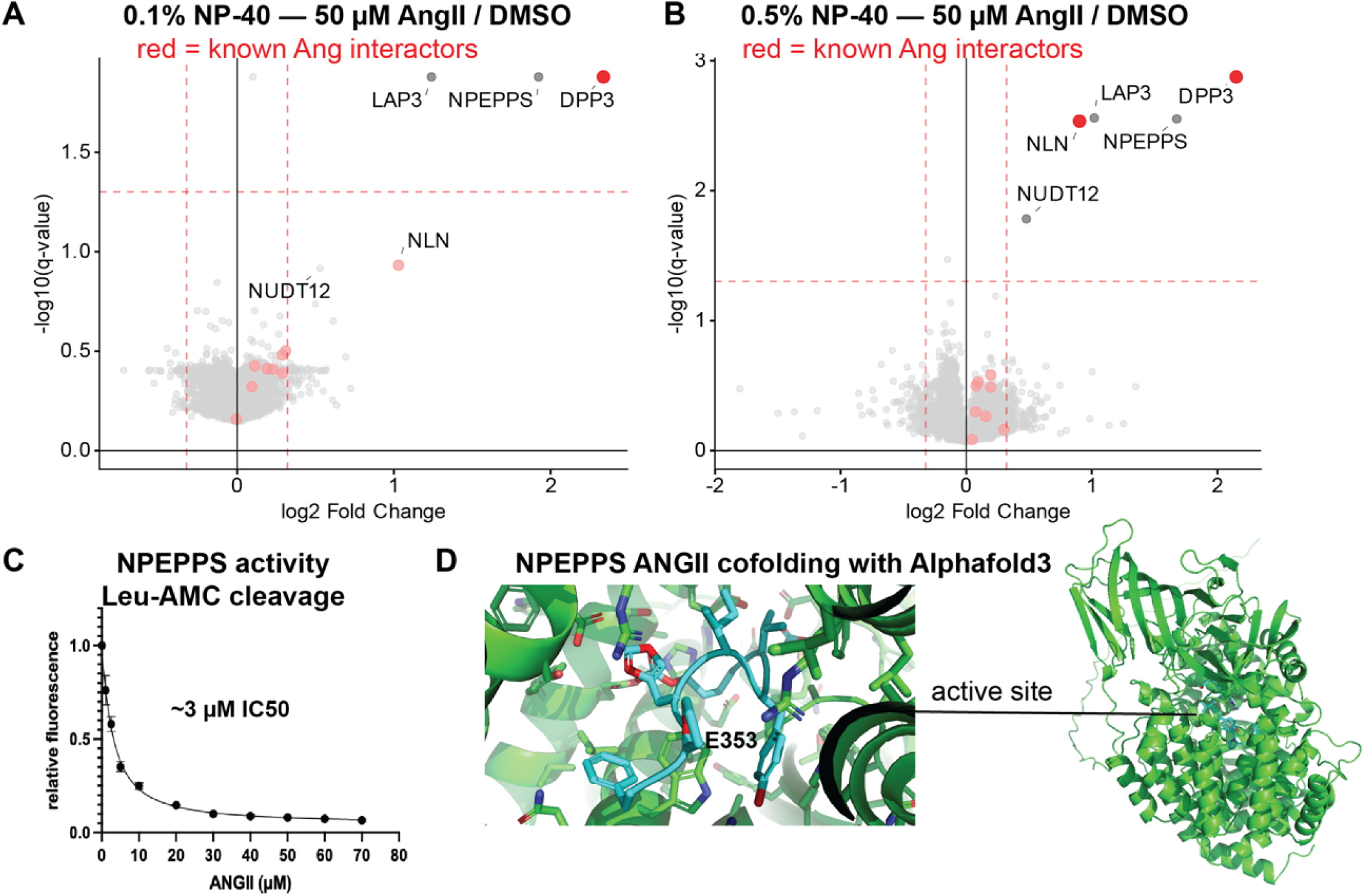
NPEPPS is a novel Angiotensin II binding protein. (A) Volcano plot showing 50 µM AngII PISA (53-59°C, centrifugation) using 0.1% NP-40 cell lysate. Known interactors for both AngI and AngII are shown in red. Lines drawn at significance thresholds of |log2fc| > 0.3 and q-value < 0.05 (B) Volcano plot showing 50 µM AngII PISA (53-59°C, centrifugation) using 0.5% NP-40 cell lysate. Known protein binders for both AngI and AngII are shown in red. Lines drawn at significance thresholds of |log2fc| > 0.3 and q-value < 0.05. (C) NPEPPS activity measured as fluorescence from the cleavage of Leu-MCA. AngII can effectively outcompete this substrate with an EC50 of 3 µM. (D) AlphaFold-3 cofolding of AngII with NPEPPS, consistent with active-site occupancy. AngII is placed near residue E353 which has been shown to reduce enzymatic activity in mutagenesis studies(^40^). Cyan is AngII and green is NPEPPS.

To generate a sufficiently large hit set for comparing melting behavior across detergent conditions, we applied a permissive raw p-value threshold (p < 0.05, |log2FC| > 0.3) rather than our standard BH-corrected threshold, detecting 126 total interactors with a small overlap (8 proteins) between conditions (**Supplementary Figure 4A**). Of the AngII hits detected exclusively in 0.5% NP-40, 22% were non-melting proteins in the 0.1% NP-40 TPP dataset, defined as having bottom plateau values > 0.4 with a poorly fit sigmoidal curve (R² < 0.8) that precludes a reliable Tm estimate (**Supplementary Figure 4B**). For example. our TPP data show that LSM1, KMT5A, and NEK4 melt poorly under 0.1% NP-40, and APIP shows a completely flat melting profile with no detectable transition, consistent with the absence of a measurable PISA fold change for these proteins at 0.1% NP-40 (**Supplementary Figure 4C, D**).This suggests that at 0.1% NP-40, these proteins undergo insufficient denaturation to produce a measurable fold change, and that higher detergent concentration enables detection of this group by lowering their Tm and shifting their melting transition into the temperature range used in the experiment. Notably, although APIP shows no melting at 0.1% NP-40, its partial denaturation under 0.5% NP-40 is sufficient to produce a measurable relative change, generating a unique hit (**Supplementary Figure 4C, D**).

We prioritized for validation the interactions that passed the q-value threshold and were reproducible across both detergent conditions, reasoning that interactions robust to a change in lysis condition are the most likely to reflect genuine binding rather than condition-specific behavior; NPEPPS, LAP3 and DPP3 were among these reproducible hits.

### Angiotensin II binds and inhibits NPEPPS

At both detergent concentrations, puromycin-sensitive aminopeptidase (NPEPPS) exhibited the second-largest fold-change, exceeded only by the known interactor DPP3. NPEPPS is an M1-family metalloaminopeptidase that degrades polyglutamine sequences released by the proteasome and participates in neuropeptide processing and antigen presentation (^29^). While NPEPPS shares aminopeptidase activity with enzymes known to process angiotensin peptides and has been proposed to participate in the renin-angiotensin-aldosterone system (RAAS), it has not been tested for angiotensin-metabolizing activity (^34, 35^). However, zinc dependent metallopeptidases from the same M1 family, including aminopeptidase A (ENPEP) and aminopeptidase N (ANPEP), and leucyl-cystinyl aminopeptidase (LNPEP), are well-established enzymes that sequentially convert angiotensin II into downstream peptides such as angiotensin III to regulate blood pressure (^36, 37^). These observations prompted us to pursue further biochemical validation of NPEPPS as a potential novel AngII interacting protein. Using recombinant human NPEPPS, we confirmed that AngII inhibits hydrolysis of the fluorogenic substrate Leu-MCA with IC₅₀ ≈ 3 μM (**Figure 5C**). Mass spectrometry analysis of AngII incubated with NPEPPS revealed no accumulation of cleavage products, indicating binding without catalytic processing (data not shown). AlphaFold3 cofolding placed AngII within the NPEPPS active site, rationalizing the observed inhibition (**Figure 5D**).

This behavior accords with NPEPPS substrate selectivity. NPEPPS preferentially cleaves substrates bearing N-terminal basic residues(^29^). ANGII (DRVYIHPF) has an N-terminal aspartate, explaining how it occupies the active site to inhibit activity without undergoing hydrolysis. Analysis of predicted NPEPPS-AngII contacts showed greater number of contacts at the N-terminal Asp-Arg dipeptide than the C-terminal hexapeptide (**Supplementary Figure 6A-C**). NPEPPS thus represents a previously unrecognized AngII-interacting protein that could potentially be modulated by the renin-angiotensin system through substrate competition.

## DISCUSSION

Thermal shift proteomics assays provide unbiased discovery of protein-ligand interactions, yet incomplete recovery of known targets and suboptimal reproducibility across research groups (^7^) complicates interpretation. We systematically evaluated three experimental parameters and characterized their effects on the high-throughput PISA assay.

Detergent concentration shifts observed Tm proteome-wide. Reducing NP-40 from 0.5% to 0.1% increased median Tm by 1.5-2°C (**Figure 2**), consistent with and extending the observations of Reinhard et al. who reported this effect for ATP-binding proteins (^10^). This finding has practical implications: temperature ranges selected for one detergent concentration may inadequately sample the denaturation transition at alternative concentrations. This is an important consideration when adapting the assay for use with low or no detergent; higher melting temperatures are required to observe significant fold changes. The modest correlation between detergent sensitivity and structural disorder, assessed through AlphaFold pLDDT scores and intramolecular contact density, suggests that less structured proteins (i.e., with more flexible or disordered domains) are most susceptible to detergent-mediated destabilization (**Figure 3; Supplementary Figure 2**). Additional descriptors such as radius of gyration, mean B-factor, and fraction of aromatic residues are not independent of pLDDT, and are therefore expected to track the pLDDT-based signal rather than to define an orthogonal determinant, with no single feature explaining the majority of the variance (**Supplementary Figure 3**). This detergent sensitivity makes disordered proteins strong candidates for condition optimization: in focused applications of thermal shift assays, tuning detergent conditions could substantially increase the observable fold change for such targets.

Aggregate removal strategy affects both curve quality and Tm distributions (**Figure 1**). Filtration yielded more well-fitted sigmoidal melting curves than centrifugation, benefiting from lower plateau values in filtration-derived curves, higher proteome depth and a more uniform Tm distribution proteome wide. The effect is particularly pronounced for DNA-binding proteins. Benzonase digestion of DNA restored well-fit melting curves for affected DNA-binding proteins under centrifugation (e.g., MACROH2A1; **Supplementary Figure 1D, E, G**), indicating that bound nucleic acid shapes their apparent aggregation and solubility behavior (^38^). We observe this change in solubility behavior in both directions after removal of polymerized nucleotides (i.e., DNA/RNA), where the proteins that only fit curves in the absence of benzonase are enriched for proteins that bind RNA (ribosome structural components) (**Supplementary Figure 1F, G)**. The underlying mechanism warrants further investigation. However, in our hands, filtration can introduce substantial inter-replicate variability in PISA experiments, likely due to inconsistency in filter plate manufacturing and a higher aggregate load when temperature points are pooled, compared to the separate filtering of individual temperature points in TPP. This is particularly problematic for these assays which rely on low variance between samples to resolve small fold changes. Any improvement in aggregate removal can easily be offset by poorer replicate reproducibility. This disadvantage is exacerbated in high-throughput applications, where it is impractical to check filtering efficiency per well. Improved filtration protocols may mitigate this limitation, which would be of significant importance when trying to detect ligand binding with DNA-binding proteins.

Temperature range selection entails balance between qualitative and quantitative sensitivity (**Figure 4**). Higher ranges amplify fold-changes for some proteins but may reduce coverage by causing loss of a larger fraction of the total protein mass. The optimal range selection therefore depends on the target population: broader ranges suit discovery screens, while narrow ranges centered on known target Tm may enhance sensitivity for focused studies. In addition, if a specific protein group, such as kinases, is targeted in a PISA screening, temperature ranges can be lowered based on their Tm values under certain detergent conditions to improve quantitative sensitivity (**Supplementary Figure 5**). Considering that PISA FCs are typically small, we recommend jointly evaluating combinations of temperature ranges and detergent concentrations when trying to maximize observed fold changes.

Beyond temperature range alone, the full parameter set, including detergent concentration and aggregate removal strategy, trade off against one another, so the appropriate choice depends on the experimental goal (e.g. discovery vs targeted screens) rather than a single universal optimum.

Maximal curve quality and proteome depth favor higher detergent and filtration, whereas target recovery can require different settings. PISA reports fold changes even for proteins that lack a well-fit Tm (**Supplementary Figure 4C**), and melting temperatures differ systematically across protein classes such as kinases, membrane proteins, and DNA-binding proteins (**Supplementary Figure 5**). As a result, the parameter set that maximizes melting curve quality proteome-wide is not necessarily the one that maximizes target recovery in a particular protein class.

Our analysis is restricted to the widely used lysate-based assay, in which these three parameters are most cleanly separable. Intact-cell PISA introduces additional variables—ligand permeability, intracellular accumulation, and the post-heating lysis step—and warrants a separate systematic study, although we anticipate that the lysis-detergent effects characterized here will transfer to that setting.

Bioactive peptides regulate virtually every aspect of human physiology, yet receptors for many through which they exert their functions remain unknown, hindering mechanistic understanding and therapeutic development (^39^). There has been a lack of unbiased screening approaches to identify binding proteins for large numbers of peptide ligands. Identification of NPEPPS, among several others, as an AngII-binding protein highlights the potential of PISA for uncovering protein-peptide interactions, though we note that PISA detects physical interactions but does not establish biological activity, which requires orthogonal validation (**Figure 5**).

In summary, systematic parameter characterization enhances the design of PISA experiments. We recommend that investigators: (1) characterize Tm distributions under intended lysis conditions prior to temperature range selection and (2) jointly evaluate detergent concentrations and temperature range selection for specific protein targets. Beyond providing a framework for rational experiment design and parameter selection, this work extends the application of PISA by enabling discovery of peptide-protein interactions, as demonstrated by our identification of NPEPPS as a novel angiotensin II-binding protein whose enzymatic activity is inhibited upon engagement.

## DATA AVAILABILITY

The mass spectrometry proteomics data have been deposited to the MassIVE Archive with the data set identifier MSV000100745.

## ACKNOWLEDGMENTS

This work was funded in part by the National Institutes of Health grant CA282268 (Q. Y.).

## SUPPLEMENTAL INFORMATION

Figure S1. Aggregate removal method effect on melting curve parameters and the effect of DNA digestion on observed melting temperatures

Figure S2. Structural features correlate with detergent sensitivity

Figure S3. Expanded structural-feature analysis of detergent sensitivity across melting and non-melting proteins

Figure S4. Angiotensin II PISA shows that 0.5% NP-40 captures more non-melting hits with a 53-59°C melting range

Figure S5. Tm and detergent sensitivity are protein class dependent Figure S6. AngII-NPEPPS interaction

Table S1 protein quant data for AngII PISA experiment. Protein and peptide quant data for detergent TPP experiments for use with TPP R package.

## References

(1) Niesen, F. H.; Berglund, H.; Vedadi, M. The Use of Differential Scanning Fluorimetry to Detect Ligand Interactions That Promote Protein Stability. Nature Protocols 2007, 2 (9), 2212–2221. 10.1038/nprot.2007.321.

(2) Martinez Molina, D.; Jafari, R.; Ignatushchenko, M.; Seki, T.; Larsson, E. A.; Dan, C.; Sreekumar, L.; Cao, Y.; Nordlund, P. Monitoring Drug Target Engagement in Cells and Tissues Using the Cellular Thermal Shift Assay. Science 2013, 341 (6141), 84–87. 10.1126/science.1233606.

(3) Savitski, M. M.; Reinhard, F. B. M.; Franken, H.; Werner, T.; Savitski, M. F.; Eberhard, D.; Martinez Molina, D.; Jafari, R.; Dovega, R. B.; Klaeger, S.; Kuster, B.; Nordlund, P.; Bantscheff, M.; Drewes, G. Tracking Cancer Drugs in Living Cells by Thermal Profiling of the Proteome. Science 2014, 346 (6205), 1255784. 10.1126/science.1255784.

(4) Becher, I.; Andrés-Pons, A.; Romanov, N.; Stein, F.; Schramm, M.; Baudin, F.; Helm, D.; Kurzawa, N.; Mateus, A.; Mackmull, M.-T.; Typas, A.; Müller, C. W.; Bork, P.; Beck, M.; Savitski, M. M. Pervasive Protein Thermal Stability Variation During the Cell Cycle. Cell 2018, 173 (6), 1495– 1507.e18. 10.1016/j.cell.2018.03.053.

(5) Huang, J. X.; Lee, G.; Cavanaugh, K. E.; Chang, J. W.; Gardel, M. L.; Moellering, R. E. High Throughput Discovery of Functional Protein Modifications by Hotspot Thermal Profiling. Nature Methods 2019, 16 (9), 894–901. 10.1038/s41592-019-0499-3.

(6) Gaetani, M.; Sabatier, P.; Saei, A. A.; Beusch, C. M.; Yang, Z.; Lundström, S. L.; Zubarev, R. A. Proteome Integral Solubility Alteration: A High-Throughput Proteomics Assay for Target Deconvolution. Journal of Proteome Research 2019, 18 (11), 4027–4037. 10.1021/acs.jproteome.9b00500.

(7) Batth, T. S.; Locard-Paulet, M.; Doncheva, N. T.; Lopez Mendez, B.; Jensen, L. J.; Olsen, J. V. Streamlined Analysis of Drug Targets by Proteome Integral Solubility Alteration Indicates Organ-Specific Engagement. Nature Communications 2024, 15 (1), 8923. 10.1038/s41467-024-53240-2.

(8) Van Vranken, J. G.; Li, J.; Mintseris, J.; Wei, T.-Y.; Sniezek, C. M.; Gadzuk-Shea, M.; Gygi, S. P.; Schweppe, D. K. Large-Scale Characterization of Drug Mechanism of Action Using Proteome-Wide Thermal Shift Assays. eLife 2024, 13, RP95595. 10.7554/eLife.95595.

(9) Zhang, X.; Lytovchenko, O.; Lundström, S. L.; Zubarev, R. A.; Gaetani, M. Proteome Integral Solubility Alteration (PISA) Assay in Mammalian Cells for Deep, High-Confidence, and High-Throughput Target Deconvolution. Bio-Protocol 2022, 12 (22), e4556. 10.21769/BioProtoc.4556.

(10) Reinhard, F. B. M.; Eberhard, D.; Werner, T.; Franken, H.; Childs, D.; Doce, C.; Savitski, M. F.; Huber, W.; Bantscheff, M.; Savitski, M. M.; Drewes, G. Thermal Proteome Profiling Monitors Ligand Interactions with Cellular Membrane Proteins. Nature Methods 2015, 12 (12), 1129–1131. 10.1038/nmeth.3652.

(11) Franken, H.; Mathieson, T.; Childs, D.; Sweetman, G. M. A.; Werner, T.; Tögel, I.; Doce, C.; Gade, S.; Bantscheff, M.; Drewes, G.; Reinhard, F. B. M.; Huber, W.; Savitski, M. M. Thermal Proteome Profiling for Unbiased Identification of Direct and Indirect Drug Targets Using Multiplexed Quantitative Mass Spectrometry. Nature Protocols 2015, 10 (10), 1567–1593. 10.1038/nprot.2015.101.

(12) Li, J.; Van Vranken, J. G.; Paulo, J. A.; Huttlin, E. L.; Gygi, S. P. Selection of Heating Temperatures Improves the Sensitivity of the Proteome Integral Solubility Alteration Assay. Journal of Proteome Research 2020, 19 (5), 2159–2166. 10.1021/acs.jproteome.0c00063.

(13) Muttenthaler, M.; King, G. F.; Adams, D. J.; Alewood, P. F. Trends in Peptide Drug Discovery. Nature Reviews Drug Discovery 2021, 20 (4), 309–325. 10.1038/s41573-020-00135-8.

(14) Kuba, K.; Sato, T.; Imai, Y.; Yamaguchi, T. Apelin and Elabela/Toddler; Double Ligands for APJ/Apelin Receptor in Heart Development, Physiology, and Pathology. Peptides 2019, 111, 62–70. 10.1016/j.peptides.2018.04.011.

(15) Lau, J. L.; Dunn, M. K. Therapeutic Peptides: Historical Perspectives, Current Development Trends, and Future Directions. Bioorganic & Medicinal Chemistry 2018, 26 (10), 2700–2707. 10.1016/j.bmc.2017.06.052.

(16) Jarzab, A.; Kurzawa, N.; Hopf, T.; Moerch, M.; Zecha, J.; Leijten, N.; Bian, Y.; Musiol, E.; Maschberger, M.; Stoehr, G.; Becher, I.; Daly, C.; Samaras, P.; Mergner, J.; Spanier, B.; Angelov, A.; Werner, T.; Bantscheff, M.; Wilhelm, M.; Klingenspor, M.; Lemeer, S.; Liebl, W.; Hahne, H.; Savitski, M. M.; Kuster, B. Meltome Atlas-Thermal Proteome Stability Across the Tree of Life. Nature Methods 2020, 17 (5), 495–503. 10.1038/s41592-020-0801-4.

(17) Hughes, C. S.; Moggridge, S.; Müller, T.; Sorensen, P. H.; Morin, G. B.; Krijgsveld, J. Single-Pot, Solid-Phase-Enhanced Sample Preparation for Proteomics Experiments. Nature Protocols 2019, 14 (1), 68–85. 10.1038/s41596-018-0082-x.

(18) Navarrete-Perea, J.; Yu, Q.; Gygi, S. P.; Paulo, J. A. Streamlined Tandem Mass Tag (SL-TMT) Protocol: An Efficient Strategy for Quantitative (Phospho)proteome Profiling Using Tandem Mass Tag-Synchronous Precursor Selection-MS3. Journal of Proteome Research 2018, 17 (6), 2226–2236. 10.1021/acs.jproteome.8b00217.

(19) Zuniga, N. R.; Frost, D. C.; Kuhn, K.; Shin, M.; Whitehouse, R. L.; Wei, T.-Y.; He, Y.; Dawson, S. L.; Pike, I.; Bomgarden, R. D.; Gygi, S. P.; Paulo, J. A. Achieving a 35-Plex Tandem Mass Tag Reagent Set Through Deuterium Incorporation. Journal of Proteome Research 2024, 23 (11), 5153–5165. 10.1021/acs.jproteome.4c00668.

(20) Yu, Q.; Paulo, J. A.; Naverrete-Perea, J.; McAlister, G. C.; Canterbury, J. D.; Bailey, D. J.; Robitaille, A. M.; Huguet, R.; Zabrouskov, V.; Gygi, S. P.; Schweppe, D. K. Benchmarking the Orbitrap Tribrid Eclipse for Next Generation Multiplexed Proteomics. Analytical Chemistry 2020, 92 (9), 6478–6485. 10.1021/acs.analchem.9b05685.

(21) Eng, J. K.; Jahan, T. A.; Hoopmann, M. R. Comet: An Open-Source MS/MS Sequence Database Search Tool. Proteomics 2013, 13 (1), 22–24. 10.1002/pmic.201200439.

(22) Wu, T.; Hu, E.; Xu, S.; Chen, M.; Guo, P.; Dai, Z.; Feng, T.; Zhou, L.; Tang, W.; Zhan, L.; Fu, X.; Liu, S.; Bo, X.; Yu, G. clusterProfiler 4.0: A Universal Enrichment Tool for Interpreting Omics Data. Innovation (Cambridge (Mass.)) 2021, 2 (3), 100141. 10.1016/j.xinn.2021.100141.

(23) Cohen, J. A Power Primer. Psychological Bulletin 1992, 112 (1), 155–159. 10.1037//0033-2909.112.1.155.

(24) Jumper, J.; Evans, R.; Pritzel, A.; Green, T.; Figurnov, M.; Ronneberger, O.; Tunyasuvunakool, K.; Bates, R.; Žídek, A.; Potapenko, A.; Bridgland, A.; Meyer, C.; Kohl, S. A. A.; Ballard, A. J.; Cowie, A.; Romera-Paredes, B.; Nikolov, S.; Jain, R.; Adler, J.; Back, T.; Petersen, S.; Reiman, D.; Clancy, E.; Zielinski, M.; Steinegger, M.; Pacholska, M.; Berghammer, T.; Bodenstein, S.; Silver, D.; Vinyals, O.; Senior, A. W.; Kavukcuoglu, K.; Kohli, P.; Hassabis, D. Highly Accurate Protein Structure Prediction with AlphaFold. Nature 2021, 596 (7873), 583–589. 10.1038/s41586-021-03819-2.

(25) Grant, B. J.; Rodrigues, A. P. C.; ElSawy, K. M.; McCammon, J. A.; Caves, L. S. D. Bio3d: An R Package for the Comparative Analysis of Protein Structures. Bioinformatics 2006, 22 (21), 2695– 2696. 10.1093/bioinformatics/btl461.

(26) Tunyasuvunakool, K.; Adler, J.; Wu, Z.; Green, T.; Zielinski, M.; Žídek, A.; Bridgland, A.; Cowie, A.; Meyer, C.; Laydon, A.; Velankar, S.; Kleywegt, G. J.; Bateman, A.; Evans, R.; Pritzel, A.; Figurnov, M.; Ronneberger, O.; Bates, R.; Kohl, S. A. A.; Potapenko, A.; Ballard, A. J.; Romera-Paredes, B.; Nikolov, S.; Jain, R.; Clancy, E.; Reiman, D.; Petersen, S.; Senior, A. W.; Kavukcuoglu, K.; Birney, E.; Kohli, P.; Jumper, J.; Hassabis, D. Highly Accurate Protein Structure Prediction for the Human Proteome. Nature 2021, 596 (7873), 590–596. 10.1038/s41586-021-03828-1.

(27) Plaxco, K. W.; Simons, K. T.; Baker, D. Contact Order, Transition State Placement and the Refolding Rates of Single Domain Proteins. Journal of Molecular Biology 1998, 277 (4), 985–994. 10.1006/jmbi.1998.1645.

(28) Abramson, J.; Adler, J.; Dunger, J.; Evans, R.; Green, T.; Pritzel, A.; Ronneberger, O.; Willmore, L.; Ballard, A. J.; Bambrick, J.; Bodenstein, S. W.; Evans, D. A.; Hung, C.-C.; O’Neill, M.; Reiman, D.; Tunyasuvunakool, K.; Wu, Z.; Žemgulytė, A.; Arvaniti, E.; Beattie, C.; Bertolli, O.; Bridgland, A.; Cherepanov, A.; Congreve, M.; Cowen-Rivers, A. I.; Cowie, A.; Figurnov, M.; Fuchs, F. B.; Gladman, H.; Jain, R.; Khan, Y. A.; Low, C. M. R.; Perlin, K.; Potapenko, A.; Savy, P.; Singh, S.; Stecula, A.; Thillaisundaram, A.; Tong, C.; Yakneen, S.; Zhong, E. D.; Zielinski, M.; Žídek, A.; Bapst, V.; Kohli, P.; Jaderberg, M.; Hassabis, D.; Jumper, J. M. Accurate Structure Prediction of Biomolecular Interactions with AlphaFold 3. Nature 2024, 630 (8016), 493–500. 10.1038/s41586-024-07487-w.

(29) Bhutani, N.; Venkatraman, P.; Goldberg, A. L. Puromycin-Sensitive Aminopeptidase Is the Major Peptidase Responsible for Digesting Polyglutamine Sequences Released by Proteasomes During Protein Degradation. The EMBO journal 2007, 26 (5), 1385–1396. 10.1038/sj.emboj.7601592.

(30) Mateus, A.; Bobonis, J.; Kurzawa, N.; Stein, F.; Helm, D.; Hevler, J.; Typas, A.; Savitski, M. M. Thermal Proteome Profiling in Bacteria: Probing Protein State in Vivo. Molecular Systems Biology 2018, 14 (7), e8242. 10.15252/msb.20188242.

(31) Leuenberger, P.; Ganscha, S.; Kahraman, A.; Cappelletti, V.; Boersema, P. J.; Mering, C. von; Claassen, M.; Picotti, P. Cell-Wide Analysis of Protein Thermal Unfolding Reveals Determinants of Thermostability. Science 2017, 355 (6327), eaai7825. 10.1126/science.aai7825.

(32) Jha, S.; Taschler, U.; Domenig, O.; Poglitsch, M.; Bourgeois, B.; Pollheimer, M.; Pusch, L. M.; Malovan, G.; Frank, S.; Madl, T.; Gruber, K.; Zimmermann, R.; Macheroux, P. Dipeptidyl Peptidase 3 Modulates the Renin-Angiotensin System in Mice. The Journal of Biological Chemistry 2020, 295 (40), 13711–13723. 10.1074/jbc.RA120.014183.

(33) Wangler, N. J.; Santos, K. L.; Schadock, I.; Hagen, F. K.; Escher, E.; Bader, M.; Speth, R. C.; Karamyan, V. T. Identification of Membrane-Bound Variant of Metalloendopeptidase Neurolysin (EC 3.4.24.16) as the Non-Angiotensin Type 1 (Non-AT1), Non-AT2 Angiotensin Binding Site. The Journal of Biological Chemistry 2012, 287 (1), 114–122. 10.1074/jbc.M111.273052.

(34) Pan, Y.; Sun, X.; Mi, X.; Huang, Z.; Hsu, Y.; Hixson, J. E.; Munzy, D.; Metcalf, G.; Franceschini, N.; Tin, A.; Köttgen, A.; Francis, M.; NHLBI Trans-Omics for Precision Medicine (TOPMed) Consortium TOPMed Kidney Function Working Group; Brody, J. A.; Kestenbaum, B.; Sitlani, C. M.; Mychaleckyj, J. C.; Kramer, H.; Lange, L. A.; Guo, X.; Hwang, S.-J.; Irvin, M. R.; Smith, J. A.; Yanek, L. R.; Vaidya, D.; Chen, Y.-D. I.; Fornage, M.; Lloyd-Jones, D. M.; Hou, L.; Mathias, R. A.; Mitchell, B. D.; Peyser, P. A.; Kardia, S. L. R.; Arnett, D. K.; Correa, A.; Raffield, L. M.; Vasan, R. S.; Cupple, L. A.; Levy, D.; Kaplan, R. C.; North, K. E.; Rotter, J. I.; Kooperberg, C.; Reiner, A. P.; Psaty, B. M.; Tracy, R. P.; Gibbs, R. A.; Morrison, A. C.; Feldman, H.; Boerwinkle, E.; He, J.; Kelly, T. N.; CRIC Study Investigators. Whole-Exome Sequencing Study Identifies Four Novel Gene Loci Associated with Diabetic Kidney Disease. Human Molecular Genetics 2023, 32 (6), 1048–1060. 10.1093/hmg/ddac290.

(35) Kopf, P. G.; Park, S.-K.; Herrnreiter, A.; Krause, C.; Roques, B. P.; Campbell, W. B. Obligatory Metabolism of Angiotensin II to Angiotensin III for Zona Glomerulosa Cell-Mediated Relaxations of Bovine Adrenal Cortical Arteries. Endocrinology 2018, 159 (1), 238–247. 10.1210/en.2017-00759.

(36) Peer, W. A. The Role of Multifunctional M1 Metallopeptidases in Cell Cycle Progression. Annals of Botany 2011, 107 (7), 1171–1181. 10.1093/aob/mcq265.

(37) Fitzgibbon, W. R. The Role of Aminopeptidases in Angiotensin Peptide Processing. Current Opinion in Physiology 2026, 47, 100898. 10.1016/j.cophys.2026.100898.

(38) Tan, C. S. H.; Go, K. D.; Bisteau, X.; Dai, L.; Yong, C. H.; Prabhu, N.; Ozturk, M. B.; Lim, Y. T.; Sreekumar, L.; Lengqvist, J.; et al. Thermal Proximity Coaggregation for System-Wide Profiling of Protein Complex Dynamics in Cells. Science 2018, 359 (6380), 1170–1177. 10.1126/science.aan0346.

(39) Yosten, G. L.; Harada, C. M.; Haddock, C.; Giancotti, L. A.; Kolar, G. R.; Patel, R.; Guo, C.; Chen, Z.; Zhang, J.; Doyle, T. M.; Dickenson, A. H.; Samson, W. K.; Salvemini, D. GPR160 de-Orphanization Reveals Critical Roles in Neuropathic Pain in Rodents. The Journal of Clinical Investigation 2020, 130 (5), 2587–2592. 10.1172/JCI133270.

(40) Thompson, M. W.; Govindaswami, M.; Hersh, L. B. Mutation of Active Site Residues of the Puromycin-Sensitive Aminopeptidase: Conversion of the Enzyme into a Catalytically Inactive Binding Protein. Arch. Biochem. Biophys. 2003, 413 (2), 236–242. DOI: 10.1016/s0003-9861(03)00123-1.

