## Supplemental Figure S1-6 for "Systematic Characterization of Thermal Stability Assay Parameters and Application in Discovery of Peptide-Protein Interactions"

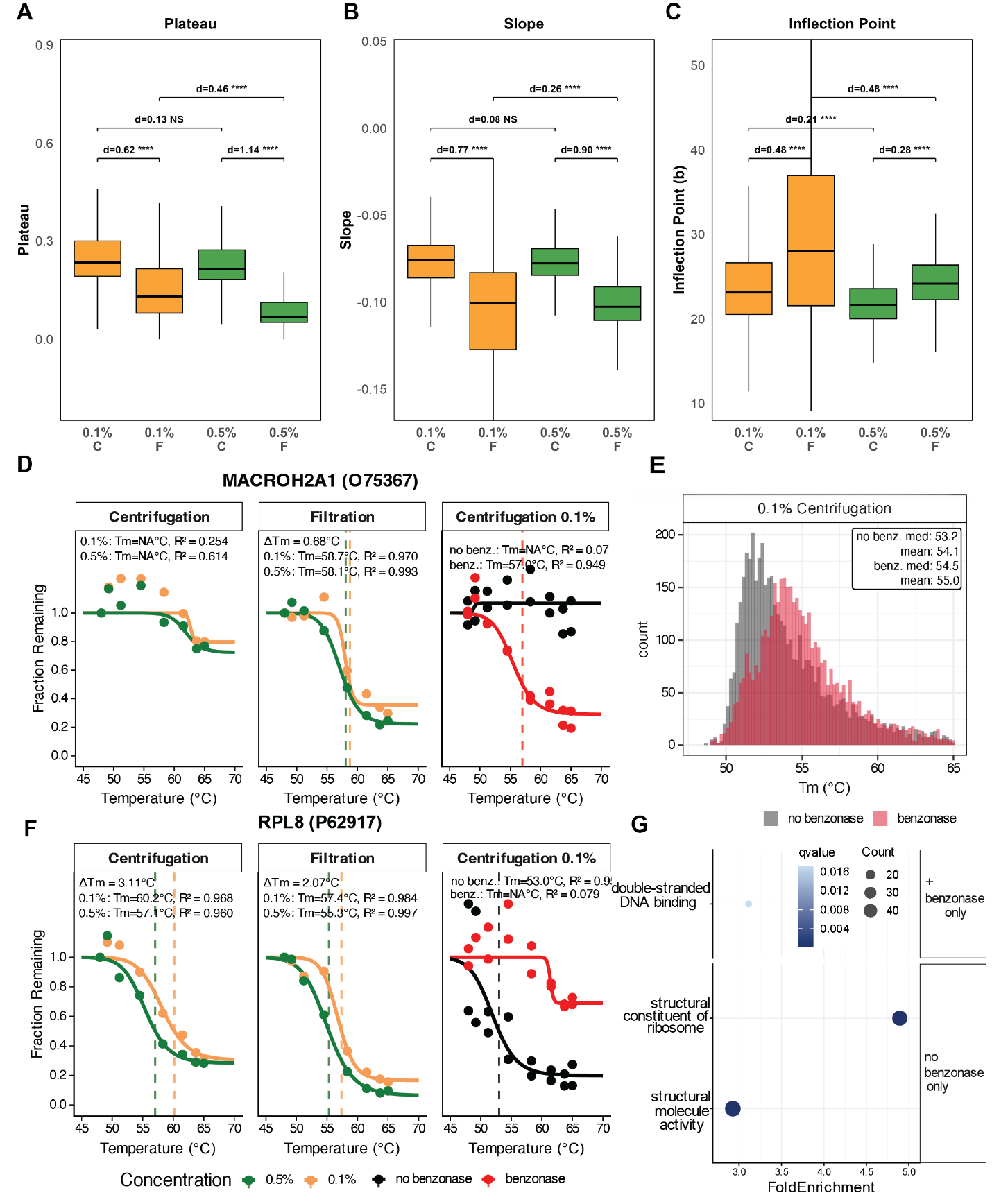

**Supplementary Figure 1. Aggregate removal method effect on melting curve parameters and the effect of benzonase digestion on observed melting temperatures**

(A) Bottom melting curve plateau distributions for both detergent concentrations (0.1% and 0.5%) and aggregate removal method (C=centrifugation, F=filtration). All proteins plotted have R^2^ > 0.95.

(B) Melting curve slope distributions for both detergent concentrations (0.1% and 0.5%) and aggregate removal method (C=centrifugation, F=filtration). All proteins plotted have R^2^ > 0.95.

(C) Inflection point (the temperature where the slope is steepest) distributions for both detergent concentrations (0.1% and 0.5%) and aggregate removal method (C=centrifugation, F=filtration). All proteins plotted have R^2^ > 0.95.

(D) Representative melting curve for DNA binding protein MACROH2A1 which fit a curve with R^2^ > 0.95 in filtration and R^2^ < 0.8 in centrifugation shows that filtration more effectively removes aggregated protein to obtain a well-fitting curve. For the same protein under 0.1% NP-40 centrifugation, benzonase pre-treatment of heated lysate (red) restored a well-fit sigmoidal melt curve relative to the untreated control (black).

(E) proteome-wide observed Tm distributions under 0.1% NP-40 centrifugation with and without benzonase pre-treatment; DNA and RNA digestion globally shifted apparent Tm (no benzonase median 53.2°C, benzonase median 54.5°C)

(F) Representative melting curve for a protein that loses its melting behavior after benzonase digestion

(G) Proteins losing a sigmoidal fit only after benzonase treatment were enriched exclusively for structural components of ribosomes. In contrast, proteins exclusively fitting sigmoidal curves in the benzonase-treated condition were enriched for double-stranded DNA-binding proteins.

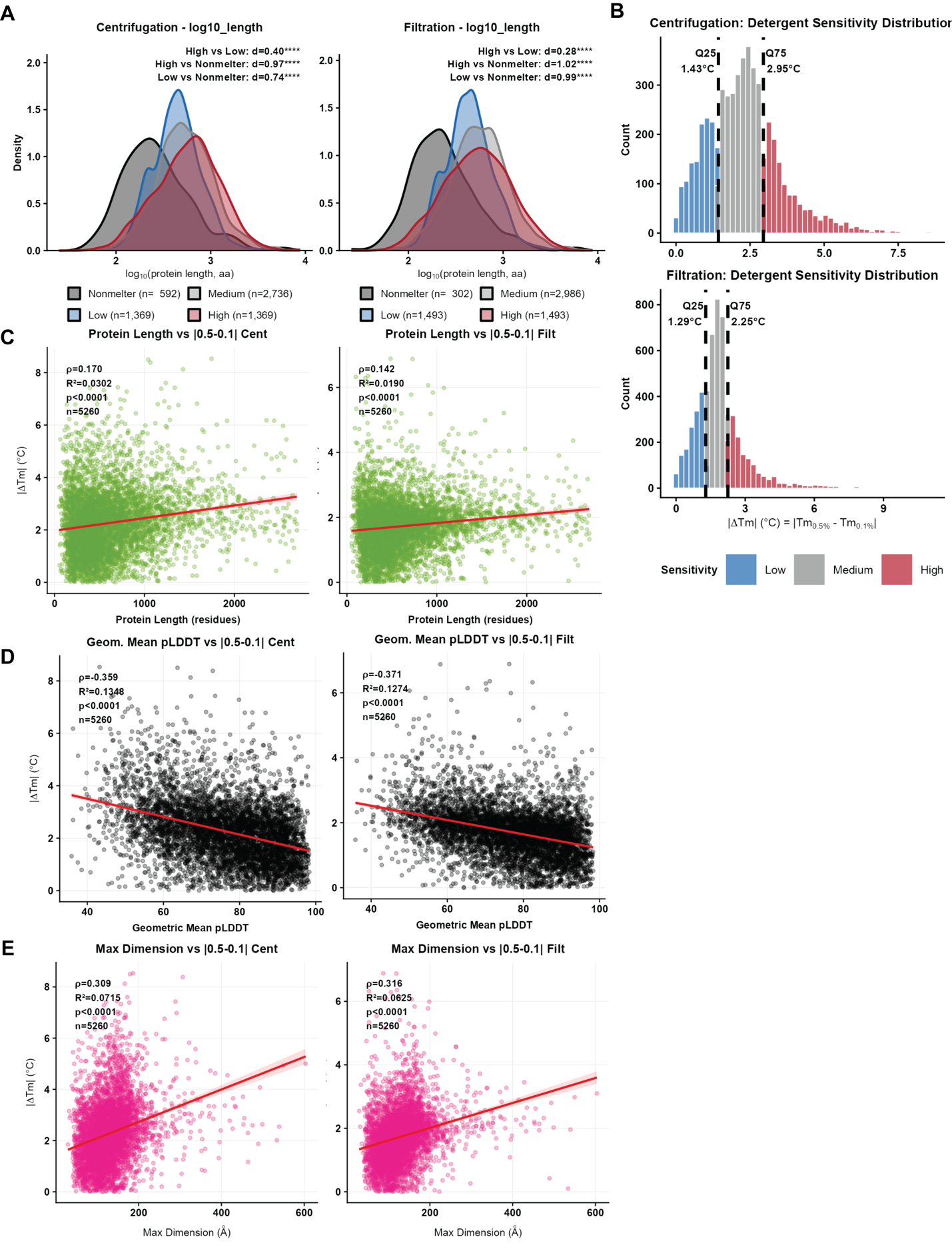

**Supplementary Figure 2. Structural features correlate with detergent sensitivity**

(A) Density of distributions of log_10_(protein length aa) for non-melting proteins and low, medium and high quartiles of detergent sensitivity for centrifugation (left) and filtration (right). Given that non-melting proteins (sigmoidal fit R^2^<0.8. plateau > 0.4) have either no or a not accurate Tm, therefore they also do not have a delta Tm value. d = [(M1 – M2) / spooled] where M1 and M2 are the group means and spooled is the pooled standard deviation represent the effect size. Effect magnitudes were interpreted using established thresholds: |d| < 0.2 negligible, < 0.5 small, < 0.8 medium, ≥ 0.8 large.

(B) Detergent sensitivity categorization into low, medium and high based on the distribution of |ΔTm| for centrifugation (top) and filtration (bottom).

(C) Spearman correlation between detergent sensitivity (|ΔTm|) and protein length with linear regression line shown in red.

(D) Spearman correlation between detergent sensitivity (|ΔTm|) and geometric mean pLDDT with linear regression line shown in red.

(E) Max dimensions (Å) of a protein assessed via pymol vs |ΔTm| (0.5% - 0.1%) for centrifugation (left) and filtration (right) methods.

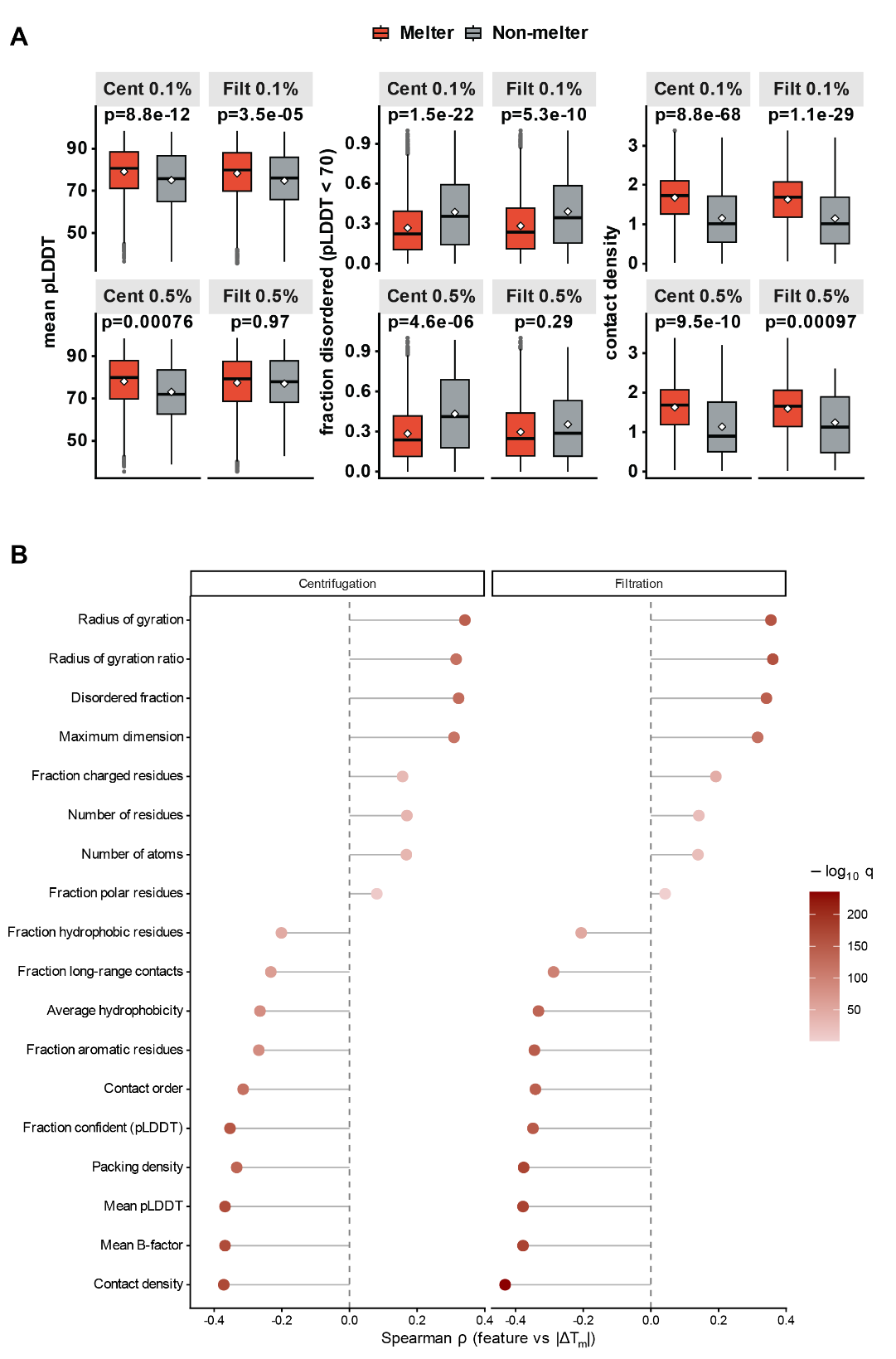

**Supplementary Figure 3. Expanded structural-feature analysis of detergent sensitivity across melting and non-melting proteins**

(A) Mean pLDDT, fraction disordered (pLDDT < 70), and intramolecular contact density for melting (red) versus non-melting (grey) proteins across all four conditions (centrifugation and filtration at 0.1% and 0.5% NP-40). White points mark medians; p-values are from two-sided Wilcoxon rank-sum tests. A protein is considered not melting if it had a plateau > 0.4 or R^2^ < 0.8.

(B) Spearman correlation (ρ) between structural features and detergent sensitivity (|ΔTm|, 0.5% NP-40 - 0.1% NP-40) for centrifugation and filtration, ranked by ρ and colored by −log10 BH-adjusted q. Features span folding confidence (mean and geometric-mean pLDDT, fraction confident), compactness (contact density, contact order, packing density, radius of gyration, max dimension), disorder (fraction disordered), and composition (length, residue and atom counts, hydrophobicity, charged/polar/aromatic fractions). No single feature explains the majority of the variance, and the strongest correlates track the pLDDT-based metrics (**Figure 3**).

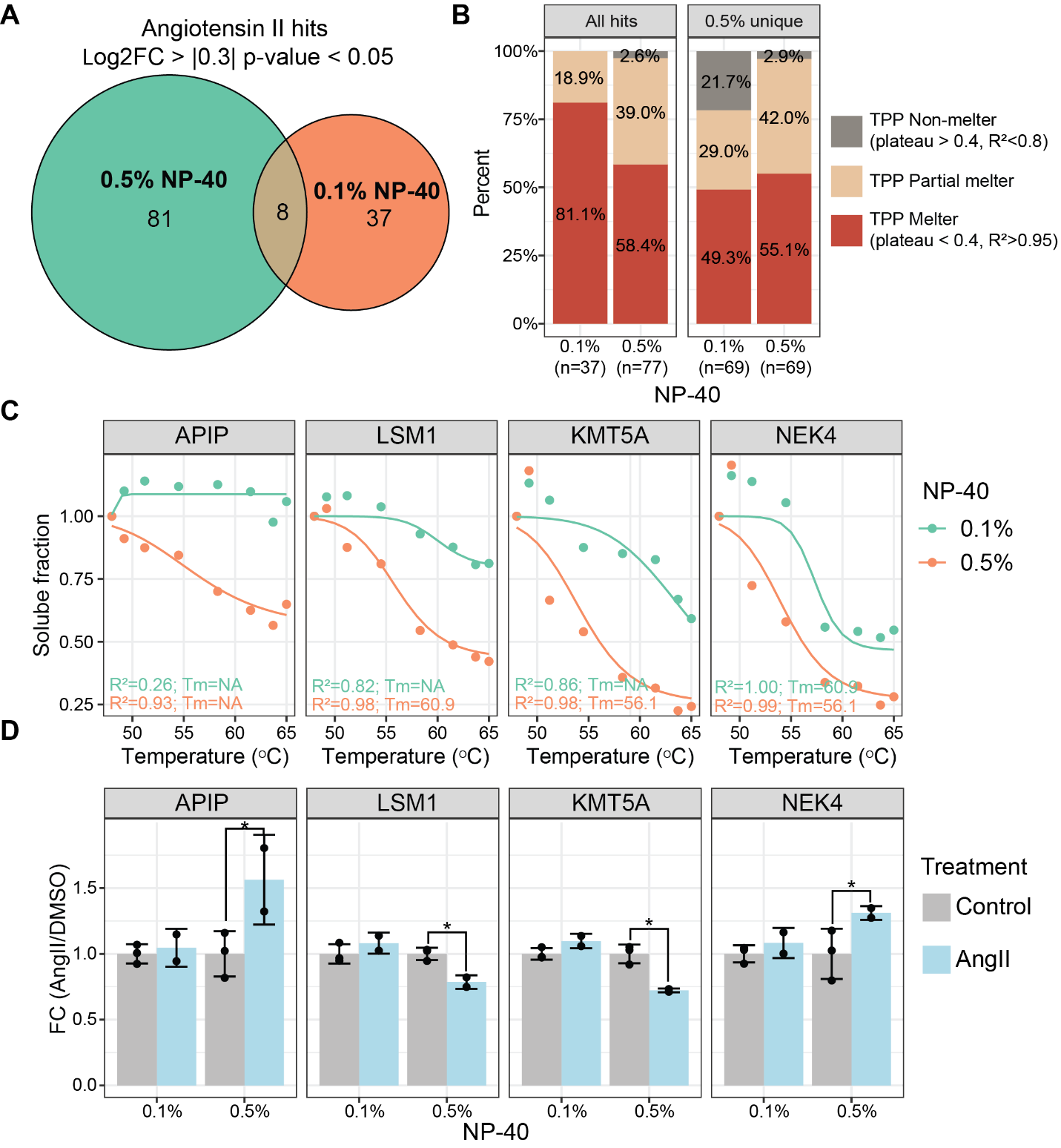

**Supplementary Figure 4. 0.5% NP-40 yields more unique Angiotensin II PISA hits, enriched for proteins that do not melt under 0.1% NP-40 and thus give no measurable PISA fold change.**

(A) Venn diagram of AngII hits passing significance thresholds (|log2FC| > 0.3, p-value < 0.05) in 0.1% and 0.5% NP-40; these supplementary hit sets use the raw p-value threshold to retain enough proteins for enrichment testing, whereas the main-text Figure 5 uses BH-adjusted q-values.

(B) Percentage of hits stratified by their TPP melting profiles: non-melter (plateau > 0.4, R²<0.8), melter (plateau < 0.4, R²>0.95), and partial melter (the remaining). Only proteins that are hits in PISA and have corresponding TPP data are kept.

(C) Representative TPP melting curves for APIP, LSM1, KMT5A, and NEK4 that were AngII PISA hits (fold change and raw p-value thresholds) uniquely under 0.5% NP-40.
(D) Corresponding PISA quantification for the same four proteins. * Student’s t-test p-value<0.05.

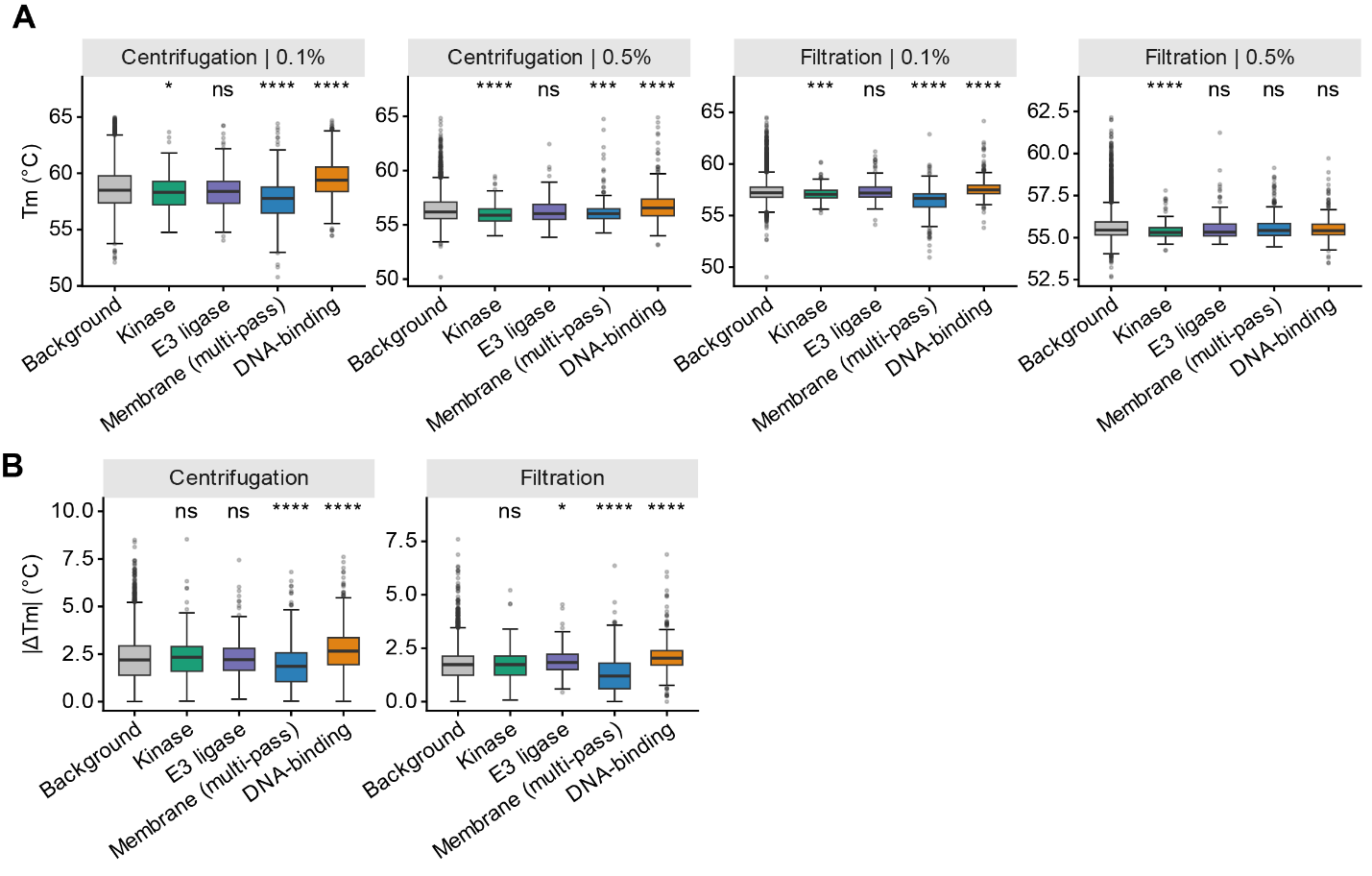

**Supplementary Figure 5. Tm and detergent sensitivity are protein class dependent.**

(A) Tm °C for 4 separate protein categories: kinases (N=221), E3 ligases (N=135), multi-pass membrane proteins (N=316) and DNA-binding proteins (N=442) against the full proteome background.

(B) |ΔTm| 0.5%-0.1% NP-40 for the same group of proteins in panel A.

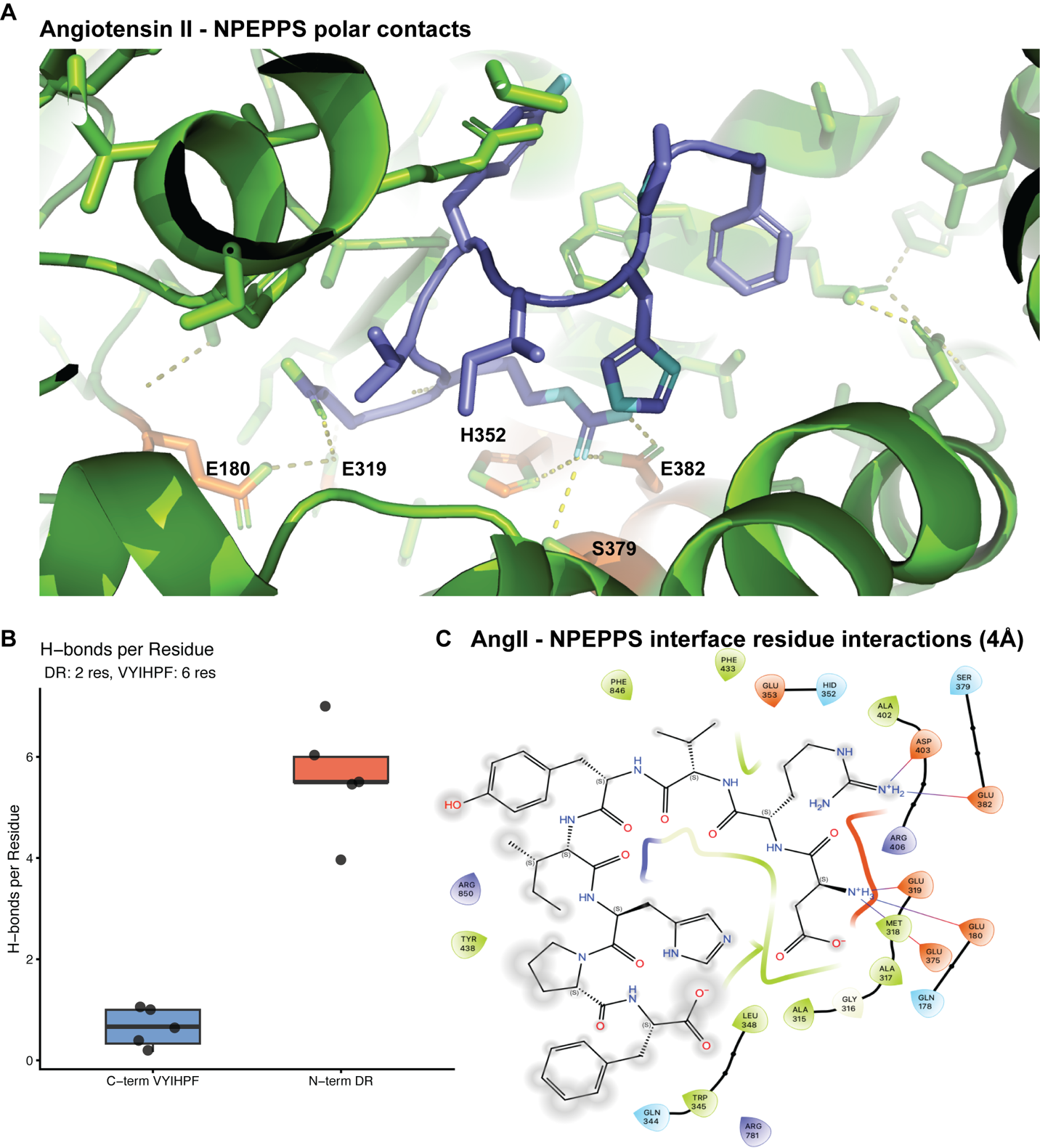

**Supplementary Figure 6. AngII-NPEPPS interaction**

(A) Visualization of interacting residues for N term DR residues versus remaining 6 amino acids using PyMol

(B) Quantification of predicted H-bonds per residue for N term DR residues versus remaining 6 amino acids using PyMol. Each dot is an individual sample within one seed.

(C) AngII interactions with the interface of NPEPPS within 4Å assessed by Maestro.
